# Identification of QTL and Underlying Genes for Root System Architecture associated with Nitrate Nutrition in Hexaploid Wheat

**DOI:** 10.1101/591529

**Authors:** Marcus Griffiths, Jonathan A. Atkinson, Laura-Jayne Gardiner, Ranjan Swarup, Michael P. Pound, Michael H. Wilson, Malcolm J. Bennett, Darren M. Wells

**Affiliations:** School of Biosciences, University of Nottingham, Sutton Bonington Campus, LE12 5RD, UK; IBM Research, Warrington, WA4 4AD, UK

**Keywords:** Doubled-haploid population, Nitrate, RNA-seq, Quantitative trait loci, Root system architecture, Triticum aestivum L. (wheat)

## Abstract

The root system architecture (RSA) of a crop has a profound effect on the uptake of nutrients and consequently the potential yield. However, little is known about the genetic basis of RSA and resource adaptive responses in wheat (*Triticum aestivum* L.). Here, a high-throughput germination paper plant phenotyping system was used to identify seedling traits in a wheat doubled haploid mapping population, Savannah × Rialto. Significant genotypic and nitrate-N treatment variation was found across the population for seedling traits with distinct trait grouping for root size-related traits and root distribution-related traits. Quantitative trait locus (QTL) analysis identified a total of 59 seedling trait QTLs. Across two nitrate treatments, 27 root QTLs were specific to the nitrate treatment. Transcriptomic analyses for one of the QTLs on chromosome 2D found under low nitrate conditions was pursued revealing gene enrichment in N-related biological processes and 17 candidate up-regulated genes with possible involvement in a root angle response. Together, these findings provide genetic insight into root system architecture and plant adaptive responses to nitrate and provide targets that could help improve N capture in wheat.

## 1 Introduction

Nitrogen (N) is an essential macronutrient for plant growth and development with agriculture greatly dependent on synthetic N fertilisers for enhancing productivity. Global demand for fertilisers is projected to rise by 1.5% each year reaching 201.7 million tonnes in 2020, over half of which (118.8 million tonnes) is for nitrate fertilizers (FAO, 2017). However, there are compelling economic and environmental reasons to reduce N fertiliser use in agriculture, particularly as the N fixing process is reliant on unsustainable fossil fuels (Dawson *et al.*, 2008).

The availability of nutrients is spatially and temporally heterogeneous in the soil (Lark *et al.*, 2004; Miller *et al.*, 2007). Roots therefore need to forage for such resources. The spatial arrangement of the root system, called the root system architecture (RSA) (Hodge *et al.*, 2009), has a profound effect on the uptake of nutrients and consequently the potential yield. Optimisation of the RSA could significantly improve the efficiency of resource acquisition and in turn increase the yield potential of the crop. An improvement in N use efficiency (NUE) by just 1% could reduce fertiliser losses and save ~$1.1 billion annually (Delogu *et al.*, 1998; Kant *et al.*, 2010).

Understanding the contribution of root traits to RSA and function is of central importance for improving crop productivity. Roots however are inherently challenging to study leading to the wide use of artificial growth systems for plant phenotyping as they are generally high-throughput, allow precise control of environmental parameters and are easy to replicate. These phenotyping systems have been key for generating root phenotypic data for association mapping and uncovering underlying genetic mechanisms (Ren *et al.*, 2012; Clark *et al.*, 2013; Atkinson *et al.*, 2015; Zurek *et al.*, 2015; Yang *et al.*, 2020). Such seedling phenotyping approaches have uncovered QTL for root system architectural traits on chromosome regions that have also been found in field trials for related traits (Bai *et al.*, 2013; Atkinson *et al.*, 2015). Only a limited number of studies have directly compared seedling screens to mature root traits in the field and overall results have been inconsistent, which likely reflects the lack of environmental control in the field, that seedling studies focus on the seminal root system and not the crown root system, and that field approaches for RSA research is in need of further development (Watt *et al.*, 2013; Rich *et al.*, 2020).

For many cereal crops, understanding the genetic basis of RSA is complex due to the polyploid nature and large genome sizes. Therefore, quantitative trait loci (QTL) analyses have been very useful for precisely linking phenotypes to regions of a chromosome. With the development of high-throughput RNA sequencing technology (RNA-seq), identified QTL can now be further dissected to the gene level. Using RNA-seq, a substantial number of genes and novel transcripts have been identified in cereal crops including rice, sorghum, maize and wheat that are implicated in RSA control (Oono *et al.*, 2013; Gelli *et al.*, 2014; Akpinar *et al.*, 2015; Yu *et al.*, 2015). To our knowledge, there are no other studies that have identified genes related to nitrate response or root angle change in wheat. The uncovering of these genes and mechanisms are likely to be of agronomical importance as they can then be implemented in genomics-assisted breeding programs to improve N-uptake efficiency in crops.

The aim of this study was to identify root traits and genes that relate to N uptake and plasticity in wheat. To achieve this, a germination paper-based system was used to phenotype a wheat doubled haploid (DH) mapping population under two N regimes. The nitrate-N levels were changed to determine the seedling responses to high- and low-affinity transport relevant concentrations as would be experienced in the field. Here were present genomic regions and underlying genes that we propose may control root size and root distribution responses in wheat to nitrate.

## 2 Results

### 2.1 Phenotypic variation in a wheat doubled haploid population for seedling traits and nitrate effects

Seedlings for 92 lines of the S×R DH mapping population and parents were grown hydroponically in a controlled environment chamber under high and low nitrate treatments (Fig. 1). Roots and shoots of each seedling were individually imaged 10 days after germination resulting in 6924 images. The results of ANOVA indicated that the variance for the genotype effects for all investigated seedling traits were highly significant (*p* < 0.001) (Table S1). Across the wheat population many of the root size and root distribution traits were found to be nitrate treatment-dependent. Interestingly, no significant differences were observed across the population for total root length in response to the nitrate treatment, however the root class distribution between lateral (*p* < 0.001) and seminal (*p* < 0.01) root length was significantly affected with a G×N-treatment interaction (*p* < 0.001). In addition, seminal root angle traits and width-depth related traits had significant nitrate treatment effects (*p <* 0.05). The seedling traits measured were also highly heritable with heritability scores for root length and count traits between 0.78-0.97, root distribution traits between 0.4-0.97, root angle traits between 0.51-0.84 and shoot traits between 0.77-0.84 (Table S1).

**Fig. 1.**
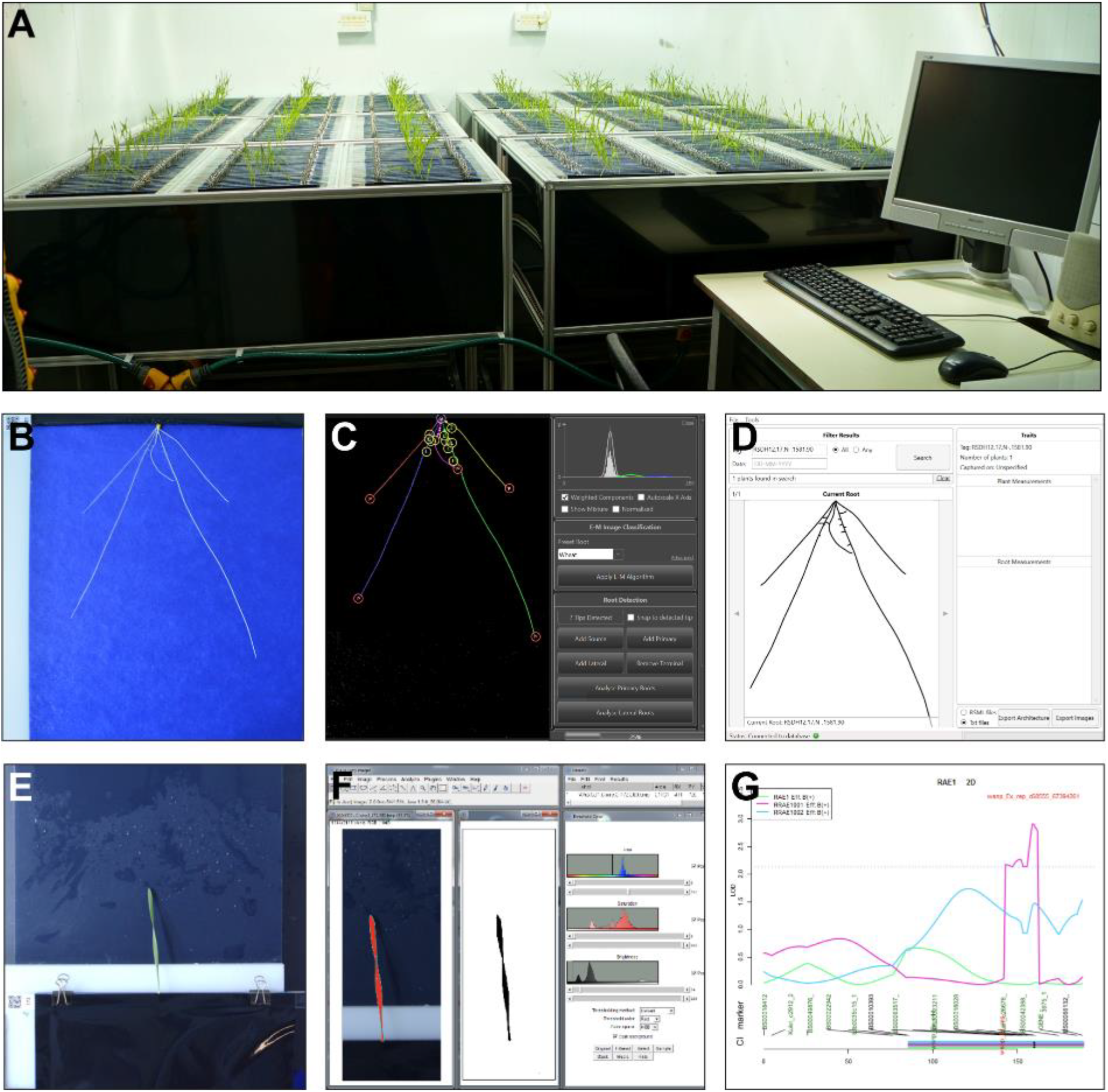
High-throughput hydroponic phenotyping system for seedling root & shoot traits. (A) Growth assembly and plant imaging station. (B) Example image of a wheat root grown on germination paper 10 days after germination. (C) Root system extraction to RSML database using RootNav software. (D) Measurement of root traits from RSML database. (E) Example image of a wheat shoot 10 days after germination. (F) Shoot image colour thresholding & shoot measurement using FIJI. (G) Example of QTL peak extracted from phenotyping data & mapping data with rQTL.

### 2.2 Wheat root phenotypic traits segregate into two distinct clusters by size and distribution

For the S×R DH population and parents a principal component analysis (PCA) was conducted to explore relationships within the root phenotypic traits (Fig. 2A). N treatment did not affect the PCA trait loadings or correlations between the traits (Fig. S1) so the analyses were conducted with both treatments together. Over 71% of the trait variation could be explained by the first two principal components and 90% of the trait variation could be explained by the first six principal components. The loadings were mostly split between root size related traits and root distribution traits (Fig. 2A). A correlation matrix of the whole dataset demonstrated the strong correlation between root size related traits and root distribution related traits (Fig. 2B). Of all the plant traits measured, the width-depth ratio traits were found to be positively correlated with the greatest number of traits from both trait groups, plant size and root distribution. In addition, the correlation analysis also highlighted negative associations between root size and angle traits.

**Fig. 2.**
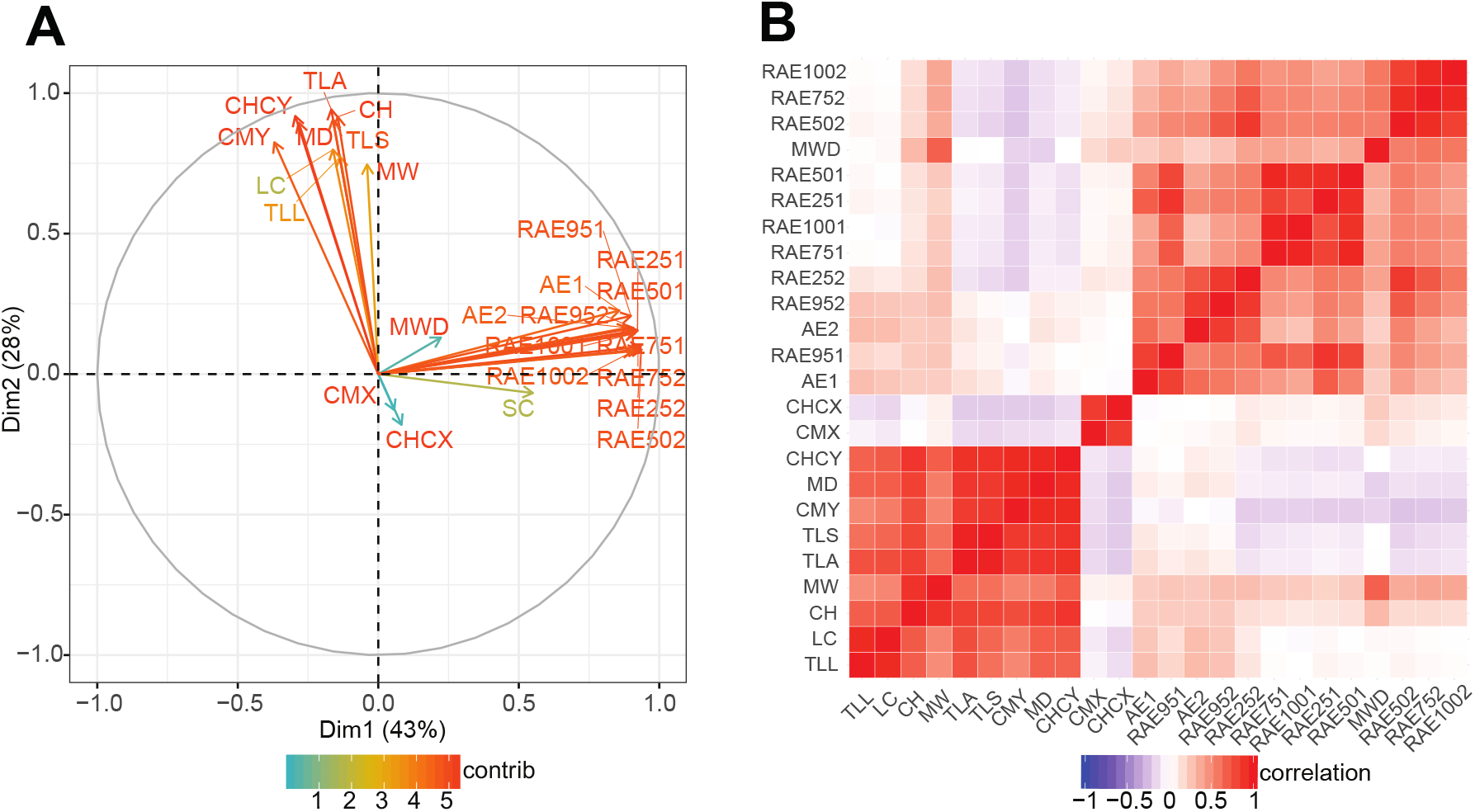
(A) PCA ordination results for S×R doubled haploid population and parents under high and low nitrate regimes. Arrows indicate directions and contributions of loadings for each trait. (B) Correlation matrix of extracted root traits averaged between nitrate treatments. Correlations are colour coded from strong positive correlation in red to strong negative correlation in blue with no correlation shown in white.

### 2.3 Identification of novel root QTLs in the S×R DH population

Using normalized phenotypic data with a high-density Savannah × Rialto iSelect map, a total of 59 QTLs were discovered for seedling traits of which 41 QTLs had positive effect alleles that came from Savannah, and 18 from Rialto (Fig. 3, Table 2). QTLs were found on chromosomes 1A, 1B, 2D, 3B, 4D, 6D, 7A and 7D, with 25 QTLs located on 6D. For the rooting traits a total of 55 QTLs were found across two nitrate treatments, 23 of which were identified under the low nitrate treatment and 32 for the high nitrate treatment. Nine root QTLs were found to be only present in the low nitrate treatment, 18 root QTLs were found only in the high nitrate treatment and 14 root QTLs (28 total) were present in both nitrate treatments. The trait ANOVA results also support the root QTLs found are nitrate condition dependent. Phenotypic variation explained by QTLs varied from 3.8 to 82.9%. Of the QTLs found, there appear to be 13 underlying root QTLs, as many root size and root distribution class traits co-localized at the same chromosome region. Two QTLs involved in shoot size traits, which were identified on chromosomes 6D and 7D under low N, were colocalized with the corresponding QTLs of root size traits. N-dependent QTLs of some traits on chromosomes 6D and 7D were colocalized with N-independent QTLs of other root size traits. For QTLs associated with nitrate treatment, QTLs for root size were found on chromosomes 1A, 6D and 7D and for root angle on chromosomes 2D, 3B and 4D. Of these regions a candidate root angle QTL (RAE1001) residing on chromosome 2D was taken forward. For this QTL, a positive allele from Rialto conferred a root angle change in the low nitrate treatment that co-localised with other root angle traits and explained 14.3% of phenotypic variation with a small peak confidence region (25 cM) (Table 2).

**Fig. 3.**
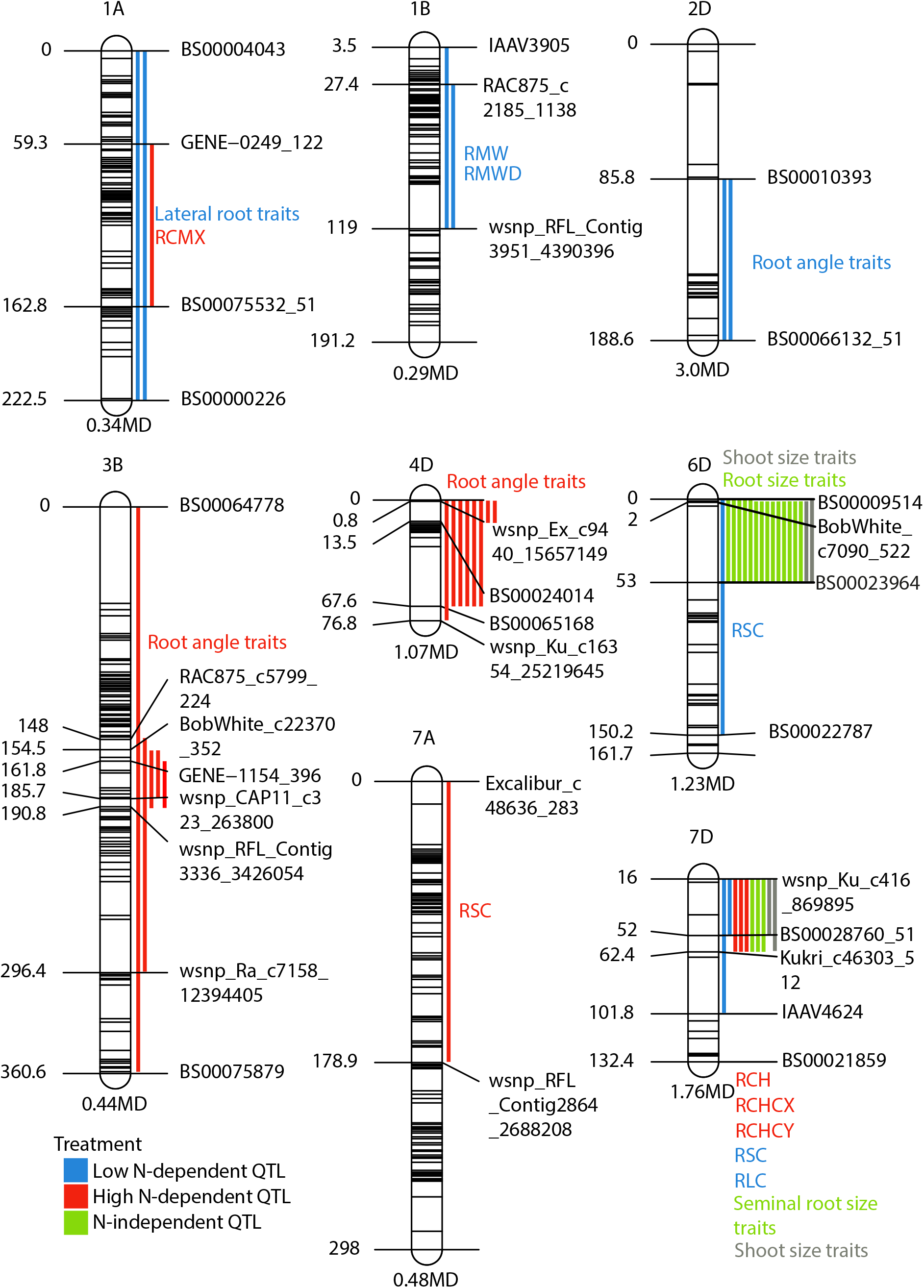
Molecular linkage map showing position of QTLs detected in the S×R DH population grown in hydroponics (LOD > 2.0). QTLs and confidence regions for all root traits are colour labelled for low N-dependent (blue), high N-dependent (red) and N treatment independent (green). Shoot QTL found in low N study are shown in grey.

### 2.4 Differentially regulated candidate genes for a root angle QTL identified by RNA-seq analysis

The lines selected for the RNA-seq analysis were based on the largest observed phenotypic differences for the trait associated with a root angle QTL located on chromosome 2D (RAE1001), found under low nitrate conditions. The DH population showed transgressive segregation with trait values more extreme than the parents (Fig. 4A). Under low nitrate there was a 30° difference in root angle (p < 0.001) between the extremes of the population with four lines of each taken forward for RNA-seq (Fig. 4B and C). The samples groups were also different for response to N with a significantly steeper root angle under low-nitrate in one of the groups (p < 0.05) (Fig. 4B).

**Fig. 4.**
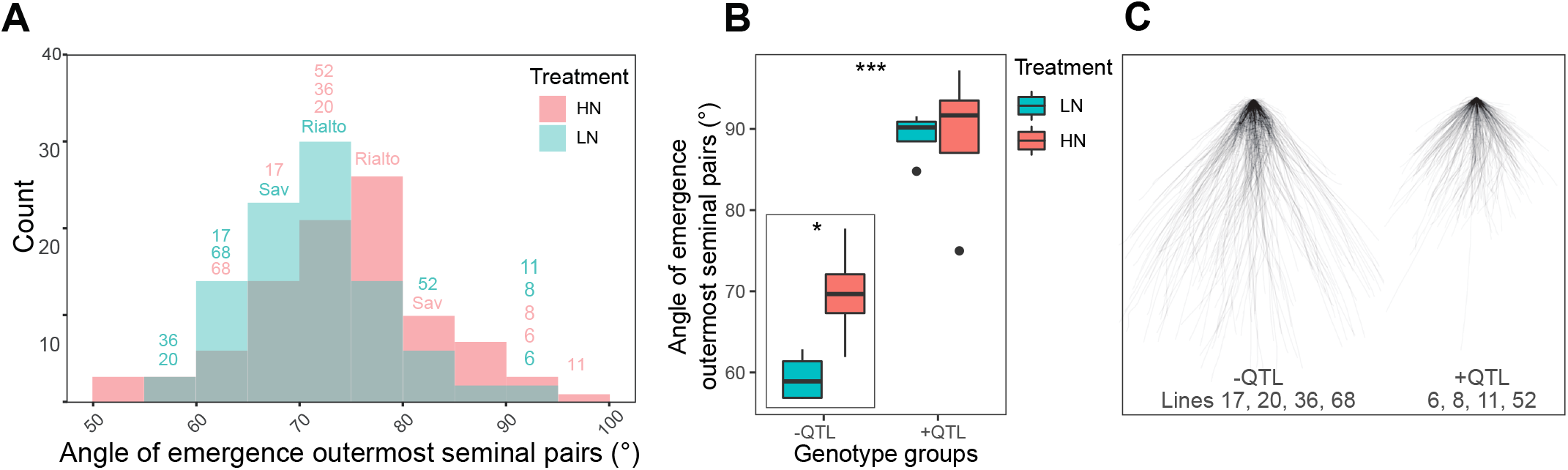
(A) Distribution of means for seminal root angle (RAE1001) for S×R doubled haploid population under two nitrate regimes. Labelled non-parental lines were selected for RNA-seq. (B) Boxplot and (C) overlay plot for lines selected for RNA-seq with differential seminal root angle (RAE1001).

One sample group was comprised of lines that had the candidate QTL with a positive effect from the parent Rialto (Group A: lines 17, 20, 36, 68) and the second sample group with parental origin from Savannah (Group B: lines 6, 8, 11, 52). As there was no single clear enriched region for the root QTL located on chromosome 2D, the whole chromosome was considered for differential gene expression analysis. A total of 3299 differentially expressed genes were identified in the analysed groups. We then focused on the identification of genes that were consistently overexpressed in Group A compared to Group B that could be driving the QTL. 1857 differentially expressed genes showed significant (*q* < 0.05) up-regulation in Group A (with the QTL) compared to Group B (without QTL) considering all four biological replicates in each case. Of these, 88 gene candidates resided on chromosome 2D. Additionally, MaSuRcA transcript assemblies were considered that were identified as significantly (*q* < 0.05) up-regulated in Group A compared Group B on chromosome 2D bringing the total to 93 (88 plus five) differentially expressed candidate sequences (Table S2). The inclusion of *de novo* assembled transcript sequences in the analysis factors for varietal specific genes that are not present in the Chinese Spring reference sequences. The sequencing read depth and alignment statistics are provided in Table S3. Of the 93 differentially expressed candidate sequences in Table S2, 17 candidate genes were consistently expressed across the Group A replicates verses zero reads mapping in one or more Group B replicates and were therefore considered as our primary candidates (Table 3). There were also 1442 differentially expressed genes that showed significant (*q* < 0.05) down-regulation in Group A (with the QTL) compared to Group B (without QTL). Of these, 65 were annotated as residing on chromosome 2D (Table S2).

Functional categories for the significantly up- and down-regulated genes were evaluated using g:profiler between contrasting sample groups for a root QTL found under low nitrate conditions. For terms relating to biological processes there were 58 up-regulated terms that had the same lowest *p*-value including “nitrogen compound metabolic process”, “cellular nitrogen compound metabolic process”, “regulation of nitrogen compound metabolic process” and “cellular nitrogen compound biosynthetic process” (Fig. 5). For the down-regulated terms, three of the top 10 terms included “nitrogen compound metabolic process”, “organonitrogen compound metabolic process” and “cellular nitrogen compound metabolic process” (Fig. 5). The complete list of enriched GO terms for molecular function, biological process and cellular component are available in Table S4. For the candidate root angle QTL found under low nitrate conditions (RAE1001) there were several N-related biological processes up- and down-regulated between the sample groups. In addition, within the candidate gene list an up-regulated NPF family gene, TraesCS2D02G348400, was identified which was consistently expressed across Group A and zero reads mapping in Group B. As this gene was expressed at low nitrate and within the identified QTL interval, *BS00010393–BS00066132_51*, the function of this gene was pursued. A phylogenetic analysis of protein families was conducted comparing NPF family protein sequences of *A. thaliana*, *O. sativa* and *T. aestivum* (Fig. S2). A total of 53 *A. thaliana* proteins, 130 *O. sativa* proteins and 391 *T. aestivum* proteins were aligned using MUSCLE with 1000 bootstrap interactions and 20 maximum likelihood searches (Edgar, 2004). The candidate *T. aestivum* protein TraesCS2D02G348400 is situated in a monocot specific sub-clade within the NPF4 clade (Fig. 6). This clade includes *A. thaliana* NPF members AtNPF4.1, AtNPF4.2, AtNPF4.3, AtNPF4.4, AtNPF4.5, AtNPF4.6 and AtNPF4.7. In addition, the candidate protein is closely related to a rice nitrate(chlorate)/proton symporter protein LOC_Os04g41410.

**Fig. 5.**
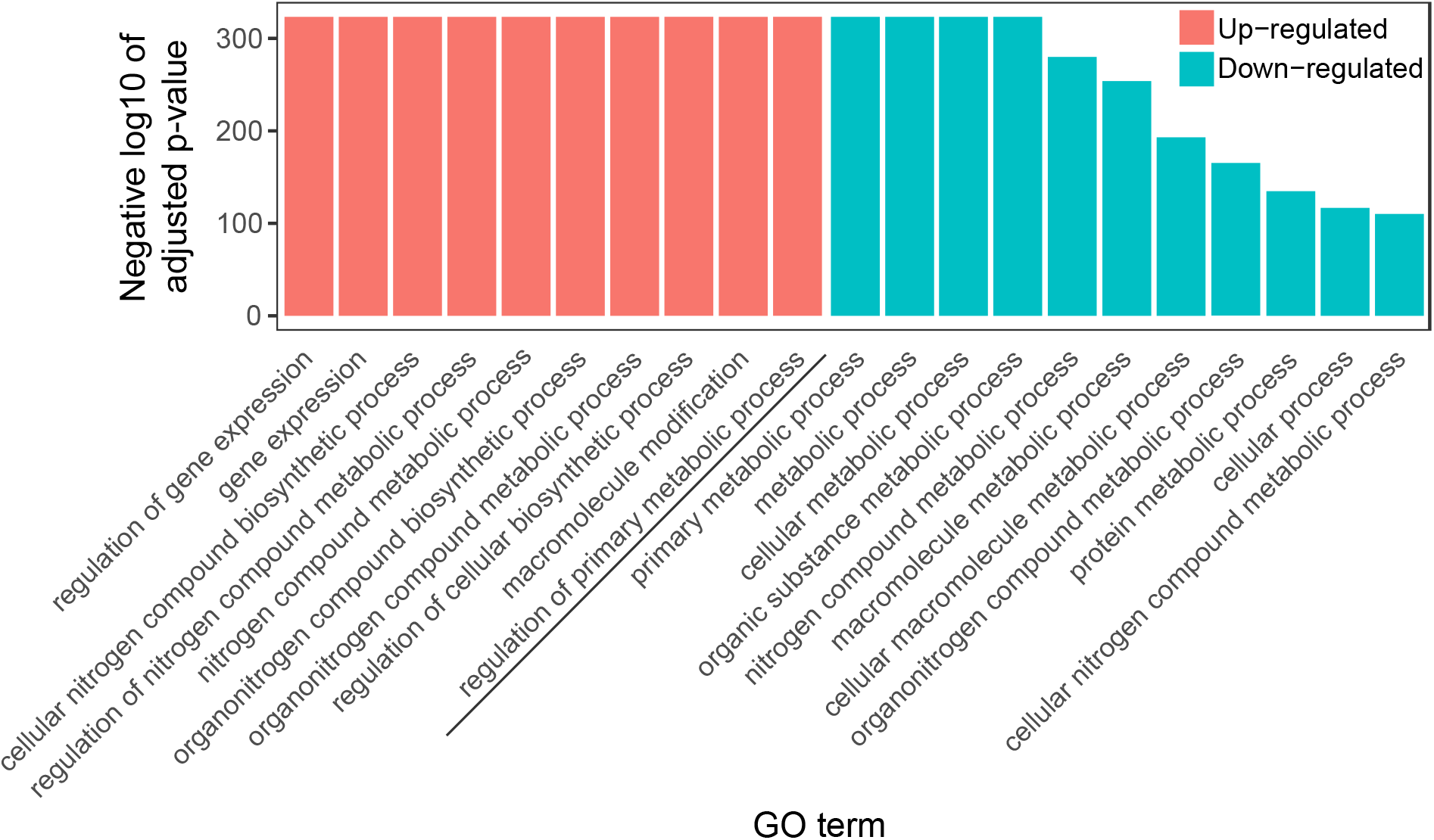
GO enrichment analysis for top Biological process GO terms with the highest p-value for up- and down-regulated genes in the sample group with a candidate seminal root angle QTL compared to without the QTL.

**Fig. 6.**
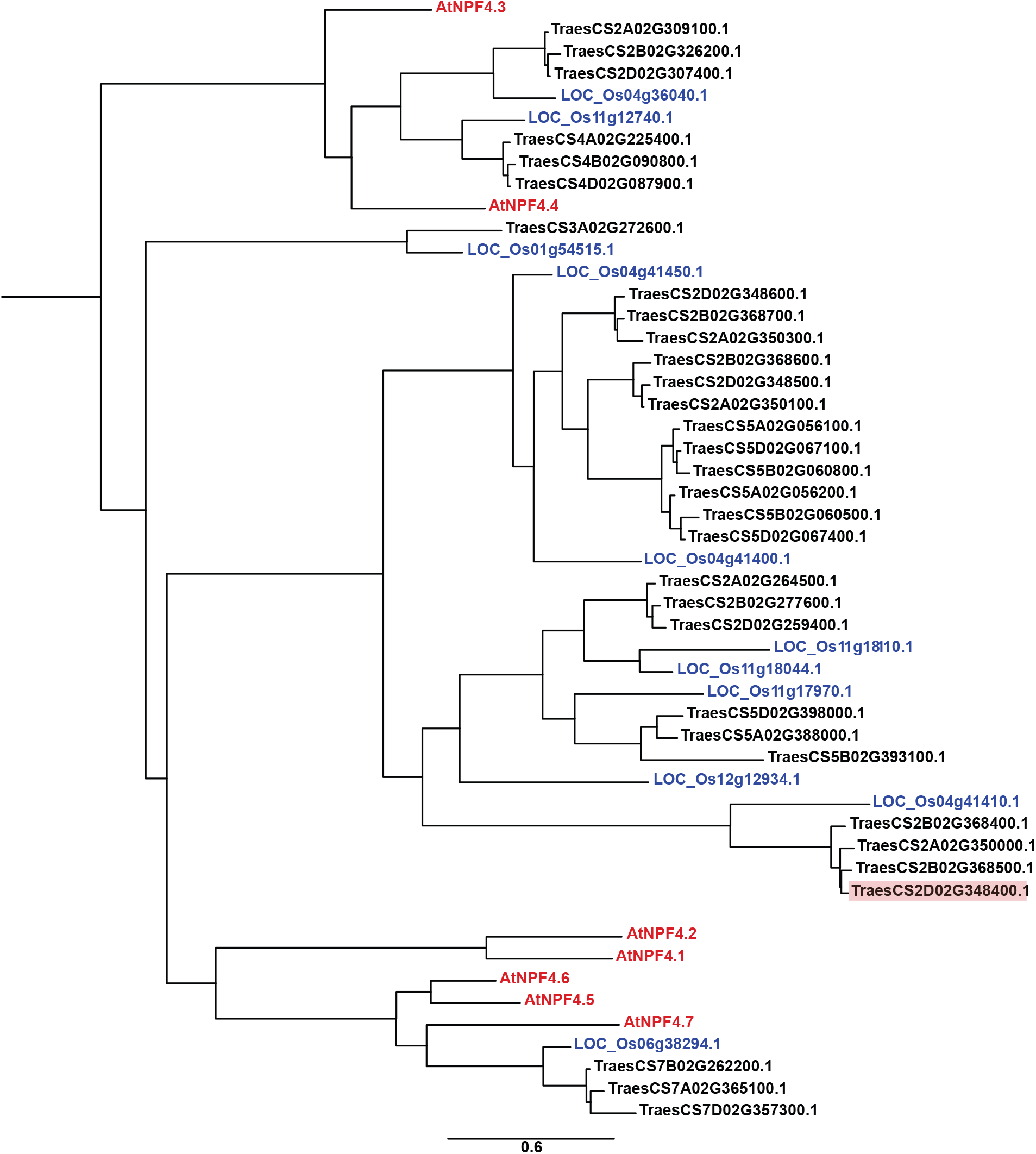
Phylogenetic tree of protein families comparing the protein sequences of A. thaliana, O. sativa L. and T. aestivum L. NPF family proteins to an identified candidate T. aestivum. protein. The candidate T. aestivum. protein is situated in a monocot specific outgroup within a NPF4 protein clade (highlighted in red). Branch lengths are proportional to substitution rate.

## 3 Discussion

Root system architecture is an important agronomic trait as the growth, development and spatial distribution of the root system affects the availability of soil resources that a plant can capture. Roots are challenging to phenotype in soil however without disturbing the spatial arrangement and therefore non-destructive root phenotyping systems such as germination paper screens and X-ray CT are invaluable tools for such work (Bai *et al.*, 2013; Atkinson *et al.*, 2015; Mooney *et al.*, 2012). In this study a germination paper-based system was used to phenotype a wheat doubled haploid (DH) mapping population under two nitrate-N regimes as it is a suitable approach for population-size root phenotyping with precise nutrition control.

Within the Savannah × Rialto DH mapping population, significant genotypic and nitrate treatment dependent phenotypic variation was observed in seedling traits (Table S1). Overall, the seedling traits could be separated into two main groups of related traits which were for root size and root distribution. In the root size trait group, it was found that the population had significant nitrate responsive plasticity in root length distribution. Interestingly the root class length of seminal and lateral roots was significantly affected by nitrate treatment, yet overall, the total root length was not significantly different. The root distribution-related traits group appeared to be the most responsive to nitrate treatment in this physiological screen using the S×R DH mapping population. Both root length distribution and root angle are widely regarded as important traits that when plastic to abiotic stimuli, such as low N or drought, as a plant can change the root foraging distribution in the soil to find such resources (Ho *et al.*, 2005; Trachsel *et al.*, 2013).

Using high-throughput methods for growth and phenotyping enabled a whole wheat DH population to be scored for root and shoot traits and the data mapped to underlying chromosome regions by QTL analysis. Nitrate plasticity is a likely a component of these QTLs where they were only detected in one of the nitrate treatments. In addition, four QTLs were found for shoot size traits on chromosomes 6D and 7D in the low nitrate conditions (LOD > 2.0) (Fig. 3, Table 2). In the literature, there are also studies that have described QTL on regions associated with those found in this study for root and shoot seedling traits. In this study on chromosome 1A, it was found that the region is associated with lateral root traits under low nitrate conditions. Interestingly, chromosome 1A has been previously associated with lateral root length in wheat and rice (Ren *et al.*, 2012; Beyer *et al.*, 2018). Therefore, there are likely underlying genes on chromosome 1A which relate to resource foraging as they were found to affect root plasticity, tolerance and/or lateral root development increase grain yield in low input agricultural systems (An *et al.*, 2006; Landjeva *et al.*, 2008; Ren *et al.*, 2012; Good *et al.*, 2017; Guo *et al.*, 2012; Zhang *et al.*, 2013; Liu *et al.*, 2013; Sun *et al.*, 2013). This chromosome 1A region has also been correlated to nitrate uptake in S×R DH field trials (Atkinson *et al.*, 2015) which would make this co-localized chromosome region an important candidate for further study with potential association between root traits and N uptake. In this study, root angle QTLs were also identified in the low nitrate conditions on chromosomes 2D, 3B and 4D. QTLs on these chromosomes have been described in other studies yet very few of these have measured root angle or distribution traits. From comparison with other studies that found root QTLs on chromosome 2D, it appears there may be an underlying gene or number of genes for seminal root development and/or plasticity (An *et al.*, 2006; Zhang *et al.*, 2013; Liu *et al.*, 2013). On chromosome 3B, other studies have found QTLs affecting root size and stress related traits or genes relating to N plasticity, uptake and mobilisation (An *et al.*, 2006; Habash *et al.*, 2007; Guo *et al.*, 2012; Zhang *et al.*, 2013; Bai *et al.*, 2013). On chromosome 4D, other studies have also found QTLs on this chromosome that indicates an underlying root development and/or root plasticity gene that may be affecting the root angle or distribution change (Zhang *et al.*, 2013; Bai *et al.*, 2013).

A seminal root angle QTL (LOD 3.0) on chromosome 2D found under low nitrate conditions was targeted for transcriptomic analysis. Significant GO enrichment in N-related biological processes were found in the chromosome region of lines with the QTL compared to those without indicating a differential low nitrate response in these lines. A total of 17 candidate up-regulated genes were identified that resided on chromosome 2D (Table 3). A more detailed list of the genes identified are given in Table S2. Two of the three genes with highest log changes plus four others have unknown function. Point mutation detection and mutant generation with TILLING or RNAi represent the next step to functionally characterise these genes. A promising candidate from the root transcriptomic analyses was an up-regulated nitrate transporter 1/peptide transporter (NPF) family gene NPF4 (TraesCS2D02G348400) in lines that had the root angle QTL. In *A. thaliana* and *O. sativa*, NPF family genes have important roles in lateral root initiation, branching and response to nitrate (Remans *et al.*, 2006; Krouk *et al.*, 2010; Fang *et al.*, 2013). However, no studies have reported genes controlling root angle change in wheat, to date. A phylogenetic analysis of protein families was conducted comparing the protein sequences of *A. thaliana*, *O. sativa* and *T. aestivum* to the candidate protein. The candidate *T. aestivum* protein is situated in a monocot specific sub-clade within the NPF4 clade and is closely related to a rice nitrate(chlorate)/proton symporter protein (LOC_Os04g41410) (Fig. 6). Members of this clade are known for transporting the plant hormone abscisic acid (ABA) (AtNPF4.1 and AtNPF4.6) and have been demonstrated to have low affinity nitrate transport activity (AtNPF4.6) (Huang *et al.*, 1999; Kanno *et al.*, 2012). ABA is known to be a key regulator in root hydrotropism, a process that senses and drives differential growth towards preferential water potential gradients (Antoni *et al.*, 2016; Takahashi *et al.*, 2002). As root angle is a determinant of root depth, pursuing this gene function is of agronomic importance for improving foraging capacity and uptake of nitrate in deep soil layers as seedling stage identified genes have been associated with yield-related traits (Xu *et al.*, 2018).

In summary, we identified 59 QTLs using a wheat seedling hydroponic system, 27 of which were for root traits found in nitrate treatment specific conditions, 14 (28 total) for root QTLs found in both treatments, and four QTLs for shoot size traits. Using transcriptome analyses we found gene enrichment in N-related biological processes which may form part of a nitrate treatment developmental response affecting root angle. These findings provide a genetic insight into plant N adaptive responses and provide targets that could help improve N capture in wheat.

## 4 Materials and methods

### 4.1 Plant materials

A winter wheat doubled haploid mapping population comprised of 94 lines was used for root phenotyping. The population was derived from an F1 plant between cultivars Savannah and Rialto (Limagrain UK Ltd, Rothwell, UK). Both parents are UK winter wheat cultivars that were on the AHDB recommended list. Savannah is a National Association of British & Irish Millers (nabim) Group 4 feed cultivar first released in 1998. Rialto is nabim Group 2 bread-making cultivar first released in 1995. Previous field research had found that Rialto had differential grain yield in low N field trials compared to Savannah making it a promising population to characterize with limited root characterization in response to N (Gaju *et al.*, 2011).

### 4.2 Seedling phenotyping

Wheat seedlings were grown hydroponically using the system described in Atkinson *et al.* (2015) (Fig. 1). Seeds from the Savannah × Rialto doubled haploid (S×R DH) mapping population were sieved to a seed size range of 2.8–3.35 mm based on mean parental seed size. Seeds were surface sterilised in 5% (v/v) sodium hypochlorite for 12 minutes before three washes in dH2O. Sterilised seeds were laid on wet germination paper (Anchor Paper Company, St Paul, MN, USA) and stratified at 4°C in a dark controlled environment room for 5 days. After stratification seeds were transferred to a controlled environment room at 20/15°C, 12-hour photoperiod, 400 μmol m^−2^ s^−1^ PAR and kept in a light-tight container. After 48 hours, uniformly germinated seedlings with ~5 mm radicle length were transferred to vertically orientated seedling pouches.

Seeds for 94 lines from the S×R DH mapping population were grown hydroponically either in high nitrate (3.13 mM NO_3_^−^, 0.75 mM NH_4_^+^) or low nitrate (0.23 mM NO_3_^−^, 0.75 mM NH_4_^+^) modified Hoagland’s solution (Table S5). The experimental design was a randomised block comprised of 94 genotypes split over 11 experimental runs with a target of 20 replications per genotype (*n* = 8 - 36). The RSA of each seedling was extracted from the images and stored in Root System Markup Language (RSML, Lobet *et al.*, 2015) using the root tracing software RootNav (Pound *et al.*, 2013). Root traits were quantified using RootNav standard functions and additional measurements as described in Atkinson *et al.* (2015). The shoot length and area were extracted from the shoot images using custom macros in the FIJI software package (Schindelin *et al.*, 2012) (macro code available in Data S1). Definitions for all extracted traits are in Table 1. Analysis of variance of the raw plant data was conducted using R package “lmerTest” (Kuznetsova *et al.*, 2017) with random effects by experimental run. Broad-sense heritability (h^2^b) was calculated using the equation 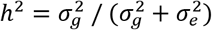 where 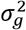 and 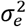 are the genetic and residual variances respectively (Falconer & Mackay, 1996). A principal component analysis (PCA) and correlation matrices were applied using R Stats Package v3.6.2 and “FactoMineR” (Husson *et al.*, 2019) using the scaled mean values to explore the relationships between the traits and genotypes within the dataset. Finally, a correlation matrix was generated using R Statistics package “corrplot” (Wei & Simko, 2017) using the raw plant data from both treatments to determine overall correlations between traits.

**Table 1.** Definition of plant traits measured.

| Acronym | Definition | Software | Units |
| --- | --- | --- | --- |
| RAE1 | Angle of emergence between the outermost seminal roots measured at 30 px | RootNav | Degrees (°) |
| RAE1001 | Angle of emergence between outermost pair of seminal roots measured at root tip | RootNav | Degrees (°) |
| RAE1002 | Angle of emergence between innermost pair of seminal roots measured at root tip | RootNav | Degrees (°) |
| RAE2 | Angle of emergence between innermost pair of seminal roots measured at 30 px | RootNav | Degrees (°) |
| RAE251 | Angle of emergence between outermost pair of seminal roots measured at first quartile of total length | RootNav | Degrees (°) |
| RAE252 | Angle of emergence between innermost pair of seminal roots measured at first quartile of total length | RootNav | Degrees (°) |
| RAE501 | Angle of emergence between outermost pair of seminal roots measured at second quartile of total length | RootNav | Degrees (°) |
| RAE502 | Angle of emergence between innermost pair of seminal roots measured at first quartile of total length | RootNav | Degrees (°) |
| RAE751 | Angle of emergence between outermost pair of seminal roots measured at third quartile of total length | RootNav | Degrees (°) |
| RAE752 | Angle of emergence between innermost pair of seminal roots measured at third quartile of total length | RootNav | Degrees (°) |
| RAE951 | Angle of emergence between outermost pair of seminal roots measured at 95 px | RootNav | Degrees (°) |
| RAE952 | Angle of emergence between innermost pair of seminal roots measured at 95 px | RootNav | Degrees (°) |
| RCH | Convex hull - area of the smallest convex polygon to enclose the root system | RootNav | mm<sup>2</sup> |
| RCHCX | Convex hull centroid - horizontal co-ordinate | RootNav | mm |
| RCHCY | Convex hull centroid - vertical co-ordinate | RootNav | mm |
| RCMX | Root centre of mass- horizontal co-ordinate | RootNav | mm |
| RCMY | Root centre of mass - vertical co-ordinate | RootNav | mm |
| RLC | Number of lateral roots | RootNav | Dimensionless (Count) |
| RMD | Maximum depth of the root system | RootNav | mm |
| RMW | Maximum width of the root system | RootNav | mm |
| RWDR | Width-depth ratio (MW/MD) | RootNav | Dimensionless (Ratio) |
| RSC | Number of seminal roots | RootNav | Dimensionless (Count) |
| RTLA | Total length of all roots | RootNav | mm |
| RTLL | Total length of lateral roots | RootNav | mm |
| RTLS | Total length of seminal roots | RootNav | mm |
| SA | Shoot area | FIJI | mm<sup>2</sup> |
| SH | Shoot height | FIJI | mm |

**Table 2.**
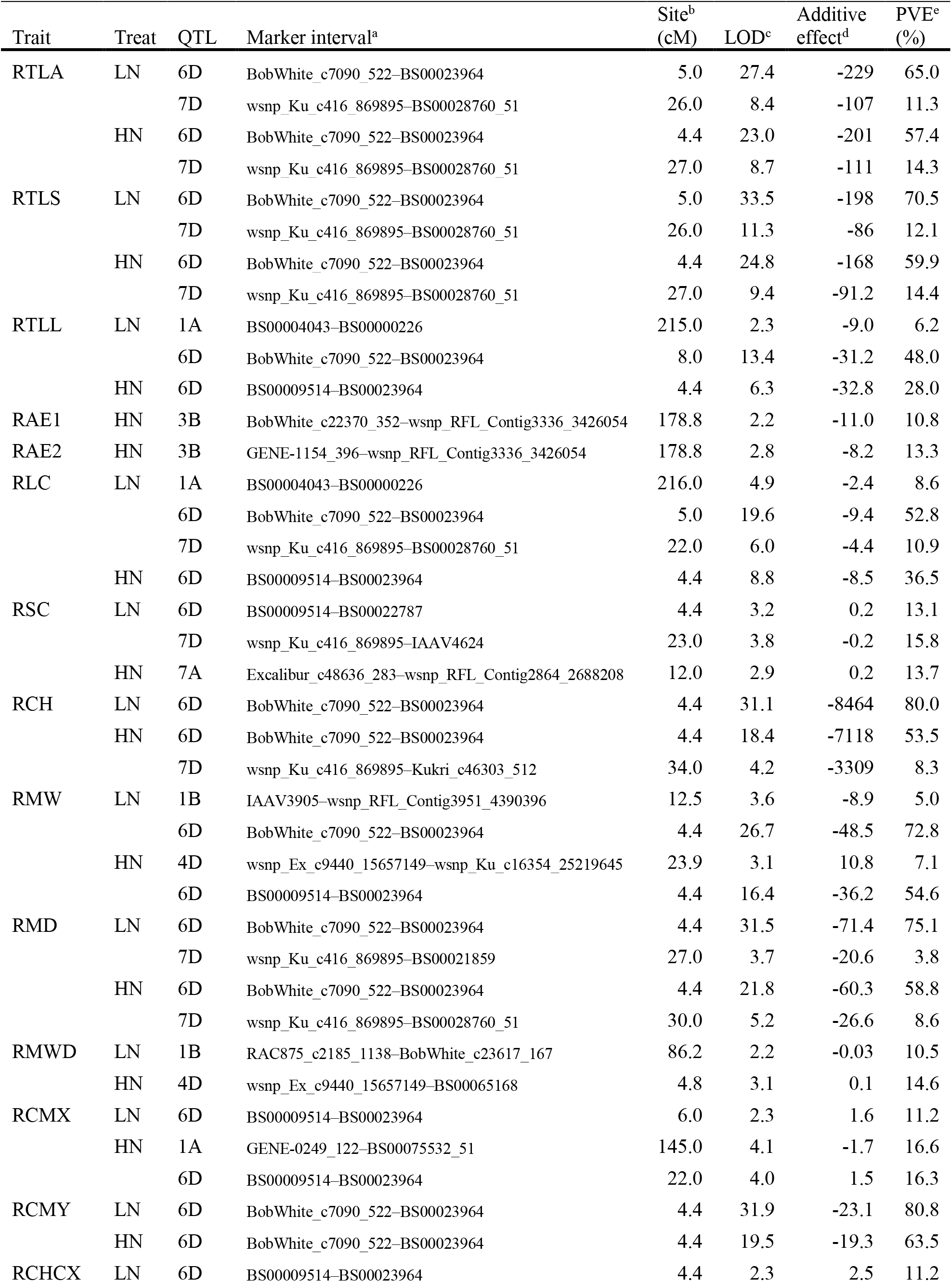
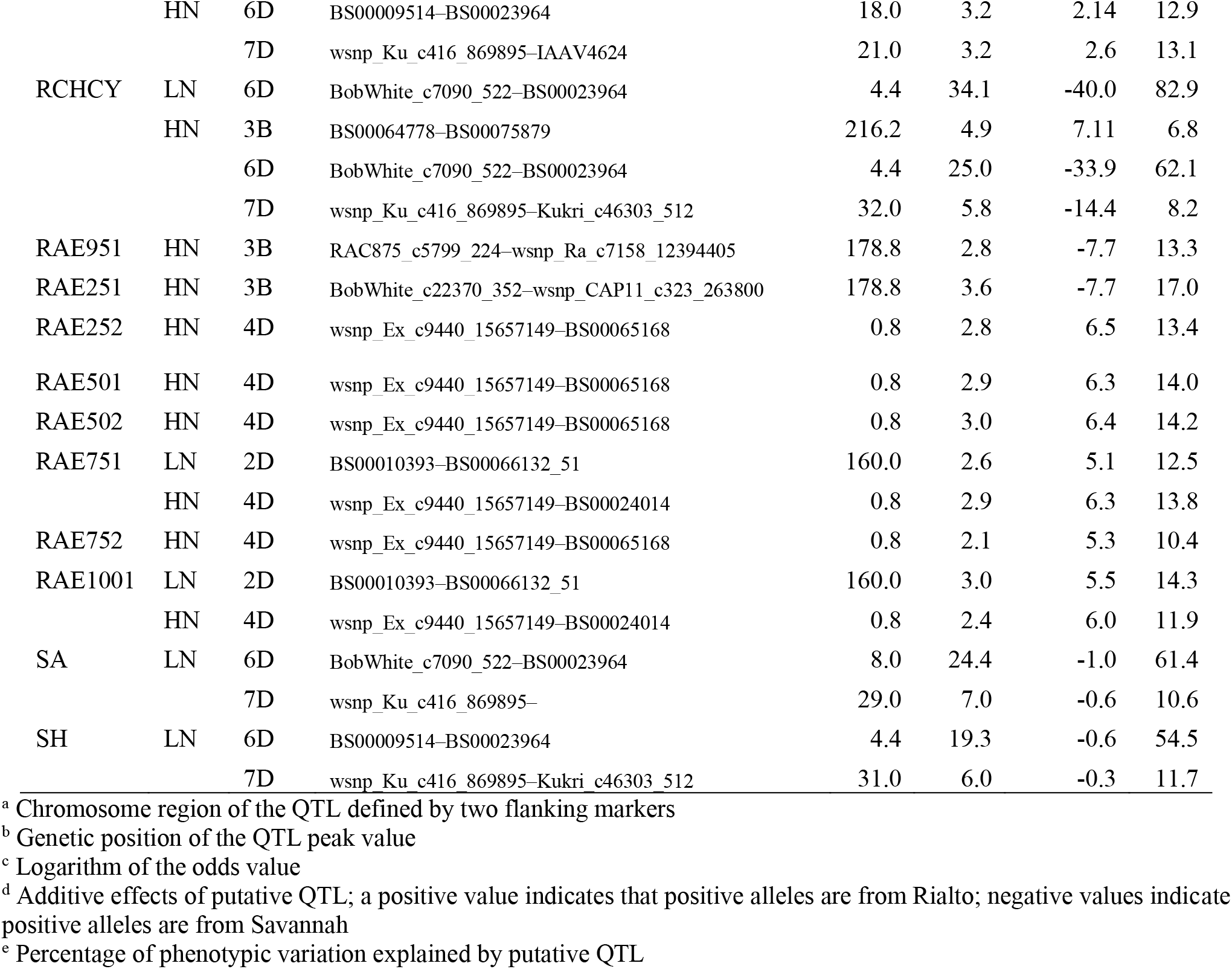
QTLs for wheat seedling traits detected in the S×R DH population grown in hydroponics (LOD > 2.0). Trait units as Table 1. Note: shoot data available for low nitrate conditions only.

**Table 3.**
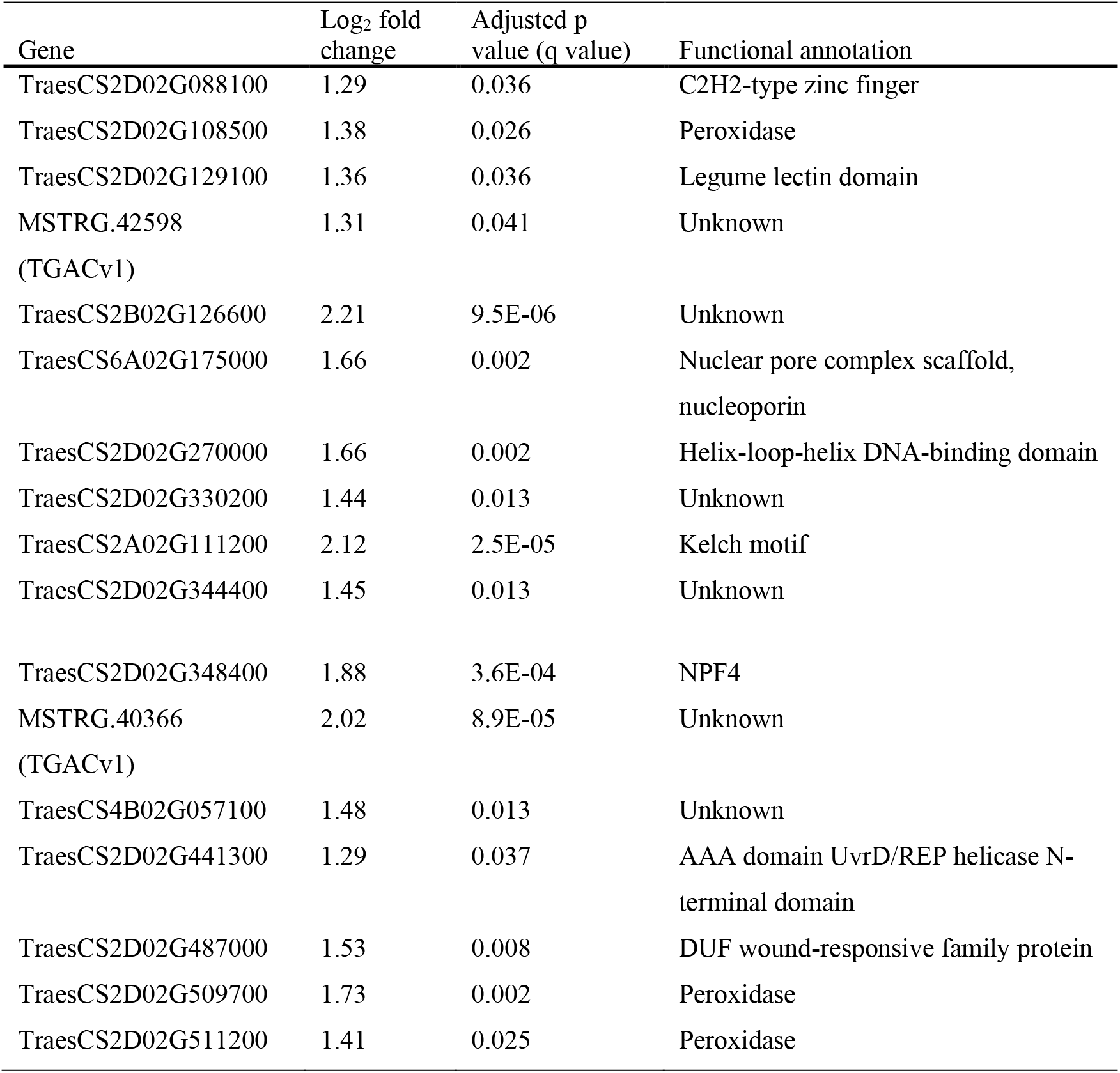
Candidate genes for seminal root angle QTL located on chromosome 2D that were consistently expressed across the Group A replicates verses zero reads mapping in one or more Group B replicates. Gene naming convention according to IWGSC RefSeq v1.1. Genes that are present on a chromosome that is not chromosome 2D represent variation between the IWGSC RefSeqv1.1 and the TGACv1 assembly.

### 4.3 Quantitative trait locus mapping

Detection of QTL and calculation of estimates for additive effects were conducted using the R Statistics package “R/qtl” (Broman *et al.*, 2003). The map used was a high-density Savannah × Rialto iSelect map obtained from Wang *et al.* (2014) with redundant and closer than 0.5 cM markers stripped out reducing the number of effective markers from 46977 to 9239 markers. Average marker density by chromosome ranged between 0.16 and 4.23 markers per cM. Before QTL analysis, best linear unbiased predictions (BLUPs) were calculated for traits showing variance between experimental runs and best linear unbiased estimations (BLUEs) calculated for all other traits (Atkinson *et al.*, 2015; Henderson, 1975; Theil, 1970) (Data S2). QTL were identified based on the composite interval mapping (CIM) via extended Haley–Knott regression (Haley & Knott, 1992). The threshold logarithm of the odds (LOD) scores and effects were calculated by 1000 × permutation test at *p* < 0.05 level (Churchill & Doerge, 1994). After the analysis, an additional threshold was applied for declaring presence of a QTL with a minimum LOD score of 2.0. The annotated linkage map was generated using R Statistics package “LinkageMapView” (Ouellette *et al.*, 2018).

### 4.4 RNA-sequencing of candidate QTL

RNA-seq was used to identify underlying genes for a candidate seminal root angle QTL (LOD 3.0) located on chromosome 2D, found under low nitrate conditions. One sample group was comprised of lines that had the candidate QTL (Group A: lines 17, 20, 36, 68) and the second sample group did not have the QTL (Group B: lines 6, 8, 11, 52). All pooled root samples of plants grown under low nitrate were collected at the same time and immediately frozen using liquid nitrogen and stored at −80°C. Each sample group had four RNA biological replicates where each replicate was a pool of roots from three plants per line (12 plants per RNA sample). Total RNA was isolated from 500–1000 mg of homogenised root tissue (TRIzol reagent). RNA quality and purity were determined using a NanoDrop™ 2000c with values above 500 ng μL^−1^ or higher accepted. Illumina 75bp Paired-End Multiplexed RNA sequencing was performed using a using NextSeq 500 by Source Bioscience (Nottingham, UK).

Differential gene expression analysis was conducted using the IWGSC RefSeq v1.1 assembly (International Wheat Genome Sequencing Consortium, 2018) (http://plants.ensembl.org/Triticum_aestivum/) and the TGAC v1 Chinese Spring reference sequence (Clavijo *et al.*, 2017). Raw sequencing reads were trimmed for adapter sequence and for regions where the average quality per base dropped below 15 (Trimmomatic version 0.32) (Bolger *et al.*, 2014). After trimming, reads below 40 bp were eliminated from the dataset. Trimmed reads were aligned to the reference sequences assembly using splice-aware aligner HISAT2 (Pertea *et al.*, 2016). Uniquely mapped reads were selected, and duplicate reads filtered out. Unmapped reads across all samples were assembled into transcripts using MaSuRCA software and sequences 250 bp or larger taken forward (Zimin *et al.*, 2013). Unmapped reads were re-aligned to these assembled transcripts individually and added to their sample specific reads while the assembled transcripts were combined with the reference sequence and GTF annotation for downstream investigations. StringTie software was used to calculate gene and transcript abundances for each sample across the analysis specific annotated genes (Pertea *et al.*, 2016). The sequencing read depth and alignment statistics are provided in Table S3. Finally, DEseq was used to visualise results and identify differential expression between samples (Anders & Huber, 2010). Differentially expressed genes were compared between the IWGSC RefSeq v1.1 and TGAC v1 reference assemblies to identify overlap using BLAST (BLASTN, e-value 1e-05, identity 95%, minimum length 40bp) (Altschul *et al.*, 1990). The top matches for each gene between the reference sequences were used to allow an integrative and comprehensive annotation of genes. Gene ontology (GO) analysis was performed with the latest genome for *T. aestivum* (IWGSC RefSeq v1.1 assembly) in g:Profiler (Reimand *et al.*, 2016) using the tailor made algorithm g:SCS for computing multiple testing correction for p-values gained from the GO enrichment analysis. A p-value threshold of 0.05 was applied with only results passing this threshold reported.

### 4.5 Phylogenetic analysis

A phylogenetic analysis of protein families was conducted to compare the protein sequences of *A. thaliana*, *O. sativa L.* and *T. aestivum L.* proton-dependent oligopeptide transporter (NPF) families (also known as the NRT1/PTR family). *A. thaliana* sequences were obtained from (Léran *et al.*, 2014). Using the latest genome for *T. aestivum* (IWGSC RefSeq v1.1 assembly) and *O. sativa* (MSU Release 7.0, Kawahara *et al.*, 2013, https://phytozome.jgi.doe.gov/) a HMM profile search was conducted (Krogh *et al.*, 2001). The resulting list of proteins were scanned using Pfam (El-Gebali *et al.*, 2019). Only single gene models of candidate genes with PTR2 domains were retained. The protein sequences were used to generate a maximum-likelihood tree using the software RAxML (Stamatakis, 2014). The exported tree file (.NWK) was then visualised using the R package “ggtree” (Yu *et al.*, 2017) and used for phylogenetic tree construction. The exported tree file (.NWK) was visualised using the R package “ggtree” (Yu *et al.*, 2017).

## 5 Data availability statement

The RNA-seq dataset is available (study PRJEB40436) from the European Nucleotide Archive (http://www.ebi.ac.uk/ena/data/view/PRJEB40436).

## 6 Funding

This work was supported by the Biotechnology and Biological Sciences Research Council [grant number BB/M001806/1, BB/L026848/1, BB/P026834/1] (MJB, DMW, and MPP); the Leverhulme Trust [grant number RPG-2016-409] (MJB and DMW); the European Research Council FUTUREROOTS Advanced Investigator grant [grant number 294729] to MG, JAA, DMW, and MJB; and the University of Nottingham Future Food Beacon of Excellence.

## 7 Disclosures

Conflicts of interest: No conflicts of interest declared

## 8 Acknowledgments

The authors would like to thank Limagrain UK Ltd for the use of the S×R DH population and Luzie U. Wingen (John Innes Centre) for providing rQTL scripts used in this work.

## Notes

### Competing Interest Statement

The authors have declared no competing interest.

### Summary of Updates

Revised text and figures.

