## Supplementary material for "Identification of QTL and Underlying Genes for Root System Architecture associated with Nitrate Nutrition in Hexaploid Wheat": Fig. S1

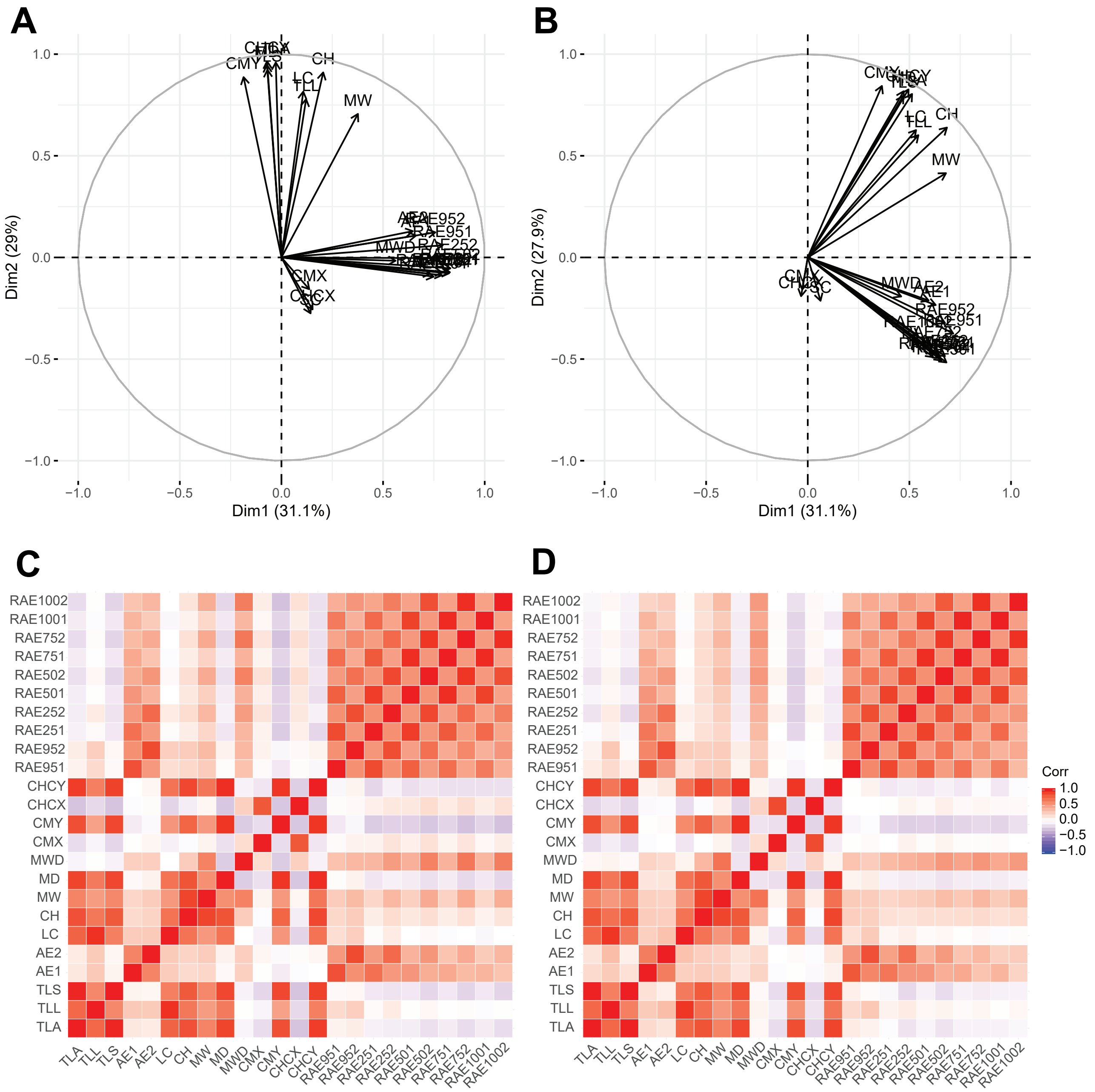


**Figure S1.** PCA ordination results for S×R doubled haploid population and parents under two N regimes, **(A)** high N and **(B)** low N. Black arrows indicate directions of loadings for each trait. Correlation matrix of extracted root traits under two N regimes, **(C)** high N and **(D)** low N. Correlations are colour coded from strong positive correlation in red to strong negative correlation in blue with no correlation shown in white.
