## Supplementary material for "Identification of QTL and Underlying Genes for Root System Architecture associated with Nitrate Nutrition in Hexaploid Wheat": Table S1

**Table S1.** Analysis of variance (ANOVA) for the investigated seedling phenotypic traits under two N regimes (n = 18, range = 8 to 36). Trait as described in Table 1. Note: shoot data available for low N treatment only.

| Trait | Source of variation | |  |  |  |
| --- | --- | --- | --- | --- | --- |
|  | Genotype | Treatment | G×N-treatment |  | Heritability (h^2^B) |
| RTLA | 39.47*** | 0.69 | 2.92*** |  | 0.97 |
| RTLS | 38.14*** | 9.4** | 2.57*** |  | 0.97 |
| RTLL | 21.55*** | 52.14*** | 5.34*** |  | 0.94 |
| RSC | 5.46*** | 4.25* | 2.05*** |  | 0.78 |
| RLC | 21.47*** | 0.69 | 4.31*** |  | 0.94 |
| RMW | 14.34*** | 12.76*** | 1.84*** |  | 0.93 |
| RMD | 42.75*** | 91.04*** | 3.42*** |  | 0.97 |
| RMWD | 3.1*** | 74.42*** | 1.45** |  | 0.62 |
| RCMX | 2.05*** | 6.51* | 1.09 |  | 0.40 |
| RCMY | 42.44*** | 2.62 | 3.72 |  | 0.97 |
| RCH | 28.36*** | 1.06 | 2.59** |  | 0.96 |
| RCHCX | 2.54*** | 0.29 | 1.14 |  | 0.54 |
| RCHCY | 45.91*** | 11.94*** | 3.48*** |  | 0.97 |
| RAE1 | 4.62*** | 1.25 | 2.97*** |  | 0.64 |
| RAE2 | 2.74*** | 8.34** | 1.72*** |  | 0.52 |
| RAE951 | 3.39*** | 43.39*** | 1.72*** |  | 0.65 |
| RAE952 | 2.34*** | 56.19*** | 1.56*** |  | 0.54 |
| RAE251 | 3.43*** | 45.19*** | 1.33* |  | 0.66 |
| RAE252 | 2.74*** | 66.13*** | 1.27* |  | 0.63 |
| RAE501 | 2.99*** | 56.52*** | 1.15 |  | 0.65 |
| RAE502 | 2.59*** | 27.24*** | 1.22 |  | 0.56 |
| RAE751 | 2.87*** | 36.73*** | 1.16 |  | 0.62 |
| RAE752 | 2.6*** | 2.09 | 1.28* |  | 0.54 |
| RAE1001 | 2.87*** | 14.84*** | 1.15 |  | 0.61 |
| RAE1002 | 2.74*** | 6.28* | 1.29* |  | 0.51 |
| SA | 5.8*** |  |  |  | 0.84 |
| SH | 3.92*** |  |  |  | 0.77 |

***denotes p<0.001; ** p<0.01; *p<0.05
