## Supplementary material for "Identification of QTL and Underlying Genes for Root System Architecture associated with Nitrate Nutrition in Hexaploid Wheat": Table S2

Table S2. Full list of up- and downregulated genes ($p < 0.05$) for a seminal roc

| Gene name | Log2 fold change | Adjusted p value | Gene start | Gene End |
|---|---|---|---|---|
| MSTRG.41571 | -1.023 | 0.027 | 13933 | 16780 |
| MSTRG.41070 | -1.421 | 0.013 | 5236 | 5836 |
| MSTRG.40319 | -4.741 | 0 | 35896 | 39333 |
| MSTRG.39723 | -5.311 | 0 | 5224 | 7444 |
| MSTRG.39722 | -1.415 | 0.013 | 65006 | 71060 |
| MSTRG.43128 | -1.761 | 0.001 | 1336 | 3353 |
| MSTRG.38813 | -2.768 | 0 | 178516 | 181072 |
| MSTRG.41475 | -1.87 | 0 | 164947 | 168405 |
| MSTRG.40665 | -0.927 | 0.016 | 36686 | 37777 |
| MSTRG.41948 | -2.79 | 0 | 81957 | 82885 |
| MSTRG.39713 | -0.776 | 0.036 | 57346 | 63900 |
| MSTRG.40021 | -4.208 | 0 | 36875 | 40738 |
| MSTRG.40926 | -1.194 | 0.012 | 31091 | 33381 |
| MSTRG.41827 | -3.046 | 0 | 98154 | 98602 |
| MSTRG.41826 | -3.73 | 0 | 94826 | 99987 |
| MSTRG.41984 | -1.422 | 0.023 | 32456 | 34869 |
| MSTRG.42351 | -1.981 | 0 | 38872 | 39198 |
| MSTRG.42167 | -2.473 | 0 | 23184 | 26021 |
| MSTRG.41606 | -5.553 | 0 | 73713 | 74667 |
| MSTRG.39361 | -1.056 | 0.049 | 85586 | 89580 |
| MSTRG.43122 | -4.627 | 0 | 1 | 721 |
| MSTRG.41746 | -1.868 | 0 | 94047 | 95805 |
| MSTRG.42880 | -1.827 | 0 | 7385 | 10104 |
| MSTRG.42988 | -0.952 | 0 | 22048 | 25876 |
| MSTRG.42825 | -3.389 | 0 | 15712 | 17589 |
| MSTRG.40551 | -1.816 | 0.001 | 15197 | 18095 |
| MSTRG.43118 | -2.278 | 0 | 2118 | 2984 |
| MSTRG.41369 | -1.896 | 0 | 148706 | 149563 |
| MSTRG.40719 | -1.327 | 0.007 | 3705 | 6272 |
| MSTRG.40222 | -3.033 | 0 | 40587 | 41386 |
| MSTRG.39288 | -2.153 | 0 | 45042 | 46459 |
| MSTRG.41543 | -2.751 | 0 | 116378 | 120735 |
| MSTRG.41546 | -6.412 | 0 | 132123 | 138655 |
| MSTRG.41371 | -1.567 | 0 | 164667 | 166875 |
| MSTRG.41372 | -2.242 | 0 | 168468 | 170738 |
| MSTRG.39299 | -4.391 | 0 | 38995 | 46253 |
| MSTRG.39298 | -2.92 | 0 | 28886 | 33783 |
| Gene.1 | -1.288 | 0.037 | 5886 | 8880 |
| MSTRG.40013 | -1.389 | 0.018 | 18869 | 29032 |
| MSTRG.42196 | -5.707 | 0 | 60267 | 65185 |
| MSTRG.41026 | -0.717 | 0.007 | 5977 | 10625 |
| MSTRG.41768 | -2.408 | 0 | 79803 | 84511 |
| MSTRG.41343 | -3.97 | 0 | 234182 | 237764 |
| MSTRG.41168 | -3.113 | 0 | 10086 | 18756 |

| | | | | |
|---|---|---|---|---|
| MSTRG.42053 | -0.773 | 0.028 | 40897 | 43290 |
| MSTRG.41064 | -0.817 | 0.043 | 20562 | 26577 |
| MSTRG.40513 | -3.217 | 0 | 16255 | 17182 |
| MSTRG.41792 | -0.828 | 0.011 | 29957 | 31740 |
| MSTRG.41676 | -2.09 | 0 | 51796 | 53092 |
| MSTRG.40221 | -1.376 | 0.022 | 63805 | 64045 |
| MSTRG.42485 | -1.426 | 0 | 14596 | 15731 |
| MSTRG.39360 | -1.312 | 0.023 | 76012 | 81026 |
| MSTRG.43144 | -1.471 | 0.012 | 1003 | 1259 |
| MSTRG.40349 | -0.799 | 0.002 | 25722 | 30634 |
| MSTRG.41333 | -1.255 | 0.044 | 159607 | 159845 |
| MSTRG.39197 | -2.227 | 0 | 64852 | 70973 |
| MSTRG.41921 | -1.434 | 0.021 | 22729 | 23007 |
| MSTRG.42740 | -1.341 | 0.015 | 14051 | 16331 |
| MSTRG.39576 | -2.161 | 0 | 84826 | 88153 |
| MSTRG.38961 | -1.49 | 0.01 | 34462 | 36245 |
| MSTRG.38889 | -0.667 | 0.002 | 75868 | 78295 |
| MSTRG.39991 | -4.686 | 0 | 6847 | 11755 |
| MSTRG.42313 | -1.12 | 0 | 26541 | 38435 |
| MSTRG.40569 | -1.415 | 0.012 | 14563 | 18117 |
| Gene.2 | -1.955 | 0 | 73310 | 75105 |
| MSTRG.42685 | 1.086453274 | 0.025534439 | 25545 | 27703 |
| MSTRG.40310 | 1.283727879 | 0.014631512 | 21996 | 23270 |
| MSTRG.39093 | 1.729993139 | 0.001733593 | 110638 | 112015 |
| TraesCS2D02G3 | 1.450005476 | 0.013427885 | 57827 | 60955 |
| TraesCS2D02G0 | 1.136254847 | 0.039296435 | 113026 | 117121 |
| MSTRG.41514 | 1.616951492 | 1.29E-05 | 79146 | 80231 |
| MSTRG.38993 | 0.949325055 | 0.032300775 | 99286 | 100318 |
| MSTRG.41956 | 1.454830842 | 0.000486757 | 14837 | 19858 |
| MSTRG.39053 | 0.974371615 | 0.000126849 | 125600 | 126408 |
| MSTRG.40097 | 1.303320656 | 2.81E-07 | 6226 | 8653 |
| MSTRG.41828 | 1.22863955 | 3.85E-10 | 33793 | 35663 |
| MSTRG.42355 | 1.319241726 | 0.036570801 | 53146 | 55510 |
| MSTRG.42598 | 1.31468 | 0.040518291 | 19073 | 19451 |
| MSTRG.42413 | 1.329835494 | 0.040325681 | 24277 | 26975 |
| MSTRG.42416 | 1.08037508 | 0.000636608 | 27131 | 30435 |
| MSTRG.39367 | 2.55799886 | 1.64E-19 | 76791 | 79575 |
| TraesCS2D02G4 | 1.28726502 | 0.036921878 | 60763 | 75025 |
| MSTRG.40726 | 2.121304672 | 2.52E-05 | 30414 | 30794 |
| MSTRG.41452 | 0.810538767 | 0.027301553 | 2996 | 4101 |
| TraesCS2D02G2 | 2.212650517 | 9.50E-06 | 5467 | 11225 |
| MSTRG.39143 | 2.983273069 | 2.36E-24 | 7939 | 9835 |
| MSTRG.39000 | 1.102970513 | 0.029790069 | 128054 | 129835 |
| MSTRG.40394 | 1.081646124 | 0.000138299 | 35587 | 36985 |
| MSTRG.42159 | 1.171985888 | 0.000449128 | 48288 | 50757 |
| MSTRG.41539 | 1.013559447 | 0.010622188 | 18927 | 21980 |

| | | | | |
|---|---|---|---|---|
| MSTRG.39328 | 0.842887972 | 0.007733659 | 13797 | 15125 |
| MSTRG.40941 | 0.808142871 | 0.008169211 | 6496 | 11297 |
| MSTRG.40576 | 1.533299371 | 0.007831497 | 20697 | 21395 |
| MSTRG.39927 | 0.940949208 | 0.02915494 | 3814 | 6280 |
| MSTRG.41621 | 1.287298951 | 0.036320236 | 34856 | 35705 |
| MSTRG.41542 | 1.06431767 | 0.040651628 | 107849 | 112998 |
| MSTRG.41775 | 1.585223259 | 6.77E-05 | 93129 | 93925 |
| MSTRG.41900 | 1.356489019 | 0.035609902 | 43987 | 46525 |
| MSTRG.40281 | 1.443305308 | 0.012862779 | 57712 | 58445 |
| MSTRG.42661 | 0.767816533 | 0.02915494 | 26103 | 28449 |
| MSTRG.40974 | 1.036676761 | 0.04723943 | 9371 | 10545 |
| MSTRG.39538 | 1.360131403 | 0.002810938 | 60816 | 62512 |
| MSTRG.39278 | 1.021496526 | 0.006551221 | 86737 | 89365 |
| MSTRG.42367 | 1.031915582 | 0.034221037 | 17477 | 19715 |
| MSTRG.38817 | 1.761440493 | 1.78E-06 | 8856 | 10980 |
| MSTRG.38818 | 1.828481699 | 4.41E-08 | 110029 | 116935 |
| MSTRG.40030 | 0.941440908 | 0.00023952 | 44788 | 46255 |
| MSTRG.39021 | 1.100988612 | 0.024344717 | 30773 | 32856 |
| MSTRG.40291 | 1.117994696 | 0.002257614 | 14091 | 16864 |
| MSTRG.41162 | 1.087705674 | 0.031728035 | 4214 | 5355 |
| MSTRG.40366 | 2.022593295 | 8.89E-05 | 50795 | 51227 |
| MSTRG.42457 | 0.734239919 | 0.043475869 | 37575 | 40165 |
| MSTRG.42452 | 0.623545907 | 0.001372379 | 3246 | 10570 |
| MSTRG.38809 | 0.901114184 | 0.003471759 | 225381 | 226914 |
| MSTRG.41174 | 1.317159197 | 0.01218422 | 7707 | 9445 |
| MSTRG.42174 | 3.828532337 | 3.67E-32 | 54386 | 55397 |
| MSTRG.38875 | 1.284124771 | 0.000265554 | 2379 | 3688 |
| MSTRG.41488 | 0.677177451 | 0.005090963 | 6537 | 11156 |
| MSTRG.41870 | 1.379580697 | 0.025822143 | 87513 | 89205 |
| MSTRG.42657 | 1.754966208 | 1.53E-06 | 6101 | 9547 |
| MSTRG.39546 | 1.19938632 | 0.025405864 | 89586 | 90416 |
| MSTRG.42075 | 0.579610858 | 0.041799164 | 47316 | 49774 |
| MSTRG.40855 | 1.388714375 | 3.64E-05 | 23964 | 28163 |
| MSTRG.40741 | 2.718075781 | 2.17E-12 | 19654 | 20525 |
| MSTRG.42220 | 2.43422701 | 4.44E-12 | 60750 | 62225 |
| MSTRG.41423 | 1.637196989 | 0.003619615 | 50006 | 51668 |
| MSTRG.40657 | 0.984349134 | 0.011372911 | 16106 | 18590 |
| MSTRG.41705 | 0.963387736 | 0.016961641 | 22637 | 25035 |
| MSTRG.41588 | 1.657236533 | 0.002127833 | 39497 | 39882 |
| MSTRG.38790 | 0.828468803 | 0.000271649 | 51329 | 52135 |
| MSTRG.38842 | 0.543780991 | 0.034385294 | 39286 | 40522 |
| MSTRG.42315 | 1.181117968 | 0.004693306 | 47016 | 55169 |
| MSTRG.40165 | 0.75973531 | 0.00845268 | 39471 | 41301 |
| MSTRG.39745 | 1.404195542 | 1.74E-05 | 21106 | 22743 |
| MSTRG.39748 | 1.655470048 | 0.001812471 | 63222 | 68255 |
| MSTRG.42854 | 0.72786411 | 0.011006493 | 12510 | 15802 |

| | | | | |
|---|---|---|---|---|
| MSTRG.40823 | 1.354453059 | 0.00158711 | 28137 | 29525 |
| MSTRG.42520 | 0.651877145 | 0.037315057 | 42093 | 43370 |
| MSTRG.39992 | 1.453963457 | 0.003510653 | 172 | 2870 |
| MSTRG.39185 | 1.413372017 | 0.025442551 | 98007 | 99465 |
| MSTRG.40245 | 1.551558663 | 6.73E-05 | 19629 | 20473 |
| MSTRG.42784 | 1.515701915 | 0.000192541 | 13967 | 14685 |
| MSTRG.42866 | 1.337599142 | 0.028215461 | 6508 | 8855 |
| MSTRG.41090 | 1.477612947 | 0.013212786 | 1436 | 6752 |
| MSTRG.40833 | 1.881446169 | 0.000358374 | 10386 | 18590 |
| MSTRG.40909 | 1.363118302 | 0.014161631 | 20796 | 21678 |
| MSTRG.42916 | 1.031386604 | 0.013427885 | 8226 | 14747 |
| MSTRG.38843 | 0.642272131 | 0.018862665 | 38758 | 41985 |
| MSTRG.41298 | 1.307245277 | 0.010234084 | 223477 | 224805 |
| MSTRG.40712 | 1.095519226 | 0.003743142 | 37158 | 38410 |
| MSTRG.40902 | 0.919026346 | 0.042310257 | 5372 | 6111 |
| MSTRG.40908 | 1.595527332 | 3.74E-06 | 14216 | 15407 |
| MSTRG.41223 | 1.032768471 | 0.044548617 | 490 | 7508 |
| MSTRG.4170 | 1.332979931 | 0.040530369 | 1 | 1136 |
| MSTRG.18948 | 1.445253528 | 0.019139272 | 1 | 650 |
| MSTRG.3022 | 1.453508492 | 0.010163954 | 1 | 462 |
| MSTRG.1448 | 0.98836053 | 0.026754906 | 1 | 538 |
| MSTRG.8673 | 1.362843004 | 0.032488435 | 1 | 335 |

ɔt angle QTL located on chromosome 2D.

| Annotation | Functional annotation |
|---|---|
| TRIAE_CS42_2DS_TGACv1_177325_AA0573540 | Cysteine-rich receptor-like protein kinase 10 |
| TRIAE_CS42_2DL_TGACv1_161611_AA0559490 | Histone superfamily protein |
| TRIAE_CS42_2DL_TGACv1_159647_AA0540950 | Protein FAR1-RELATED SEQUENCE 5 |
| TRIAE_CS42_2DL_TGACv1_158781_AA0526060 | Phenylalanine ammonia-lyase |
| TRIAE_CS42_2DL_TGACv1_158779_AA0526040 | Protein PAIR1 |
| n/a | - |
| n/a | - |
| TRIAE_CS42_2DS_TGACv1_177259_AA0570990 | Pentatricopeptide repeat-containing protein |
| TRIAE_CS42_2DL_TGACv1_160293_AA0549090 | - |
| TRIAE_CS42_2DS_TGACv1_177738_AA0583520 | AT hook motif DNA-binding |
| TRIAE_CS42_2DL_TGACv1_158763_AA0525740 | TCP transcription factor |
| TRIAE_CS42_2DL_TGACv1_159163_AA0533670 | Syntaxin-61 |
| TRIAE_CS42_2DL_TGACv1_160950_AA0555540 | Benzoate carboxyl methyltransferase |
| TRIAE_CS42_2DS_TGACv1_177573_AA0580310 | - |
| TRIAE_CS42_2DS_TGACv1_177573_AA0580310 | - |
| TRIAE_CS42_2DS_TGACv1_177783_AA0584500 | Disease resistance protein RGA2 |
| n/a | - |
| TRIAE_CS42_2DS_TGACv1_178019_AA0589290 | - |
| TRIAE_CS42_2DS_TGACv1_177364_AA0574640 | - |
| TRIAE_CS42_2DL_TGACv1_158383_AA0517050 | Yellow stripe-like transporter 11 |
| n/a | - |
| TRIAE_CS42_2DS_TGACv1_177484_AA0578360 | - |
| TRIAE_CS42_2DS_TGACv1_179278_AA0605830 | RNA binding protein |
| TRIAE_CS42_2DS_TGACv1_179693_AA0608660 | Phosphatidylinositol 3- and 4-kinase protein |
| TRIAE_CS42_2DS_TGACv1_179146_AA0604650 | Exostosin-1 |
| TRIAE_CS42_2DL_TGACv1_160071_AA0546360 | - |
| n/a | - |
| TRIAE_CS42_2DS_TGACv1_177184_AA0568110 | - |
| n/a | - |
| n/a | - |
| TRIAE_CS42_2DL_TGACv1_158317_AA0515420 | - |
| TRIAE_CS42_2DS_TGACv1_177302_AA0572750 | Disease resistance protein RPP13 |
| TRIAE_CS42_2DS_TGACv1_177302_AA0572780 | Disease resistance protein RPM1 |
| TRIAE_CS42_2DS_TGACv1_177184_AA0568150 | Endochitinase 1 |
| TRIAE_CS42_2DS_TGACv1_177184_AA0568160 | Nicotianamine synthase-like 5 protein |
| TRIAE_CS42_2DL_TGACv1_158328_AA0515710 | Smr domain-containing protein |
| TRIAE_CS42_2DL_TGACv1_158328_AA0515700 | Histone H4 |
| TRIAE_CS42_2DS_TGACv1_180584_AA0611090 | F-box domain containing protein |
| TRIAE_CS42_2DL_TGACv1_159154_AA0533500 | DNA-directed RNA polymerase subunit beta |
| n/a | - |
| TRIAE_CS42_2DL_TGACv1_161369_AA0558280 | Glutamine synthetase |
| TRIAE_CS42_2DS_TGACv1_177494_AA0578800 | Retrotransposon protein |
| n/a | - |
| TRIAE_CS42_2DL_TGACv1_162104_AA0561840 | GTP-binding nuclear protein Ran-B1 |

| | |
|---|---|
| TRIAE_CS42_2DS_TGACv1_177870_AA0586390 | Endonuclease/exonuclease/phosphatase |
| TRIAE_CS42_2DL_TGACv1_161554_AA0559210 | Disease susceptibility protein LOV1 |
| n/a | - |
| TRIAE_CS42_2DS_TGACv1_177535_AA0579540 | Glycosyltransferase |
| TRIAE_CS42_2DS_TGACv1_177420_AA0576580 | Protein translocase subunit SecA |
| n/a | - |
| TRIAE_CS42_2DS_TGACv1_178531_AA0597160 | DUF679 domain membrane protein 2 |
| TRIAE_CS42_2DL_TGACv1_158383_AA0517040 | Yellow stripe-like transporter 11 |
| n/a | - |
| TRIAE_CS42_2DL_TGACv1_159701_AA0541690 | inositol transporter 1 |
| n/a | - |
| n/a | - |
| TRIAE_CS42_2DS_TGACv1_177712_AA0582950 | - |
| TRIAE_CS42_2DS_TGACv1_178972_AA0603000 | - |
| TRIAE_CS42_2DL_TGACv1_158583_AA0522510 | Nuclease S1 |
| n/a | - |
| TRIAE_CS42_2DL_TGACv1_157983_AA0504720 | Autophagy-related protein 8C |
| TRIAE_CS42_2DL_TGACv1_159133_AA0533040 | ATP-dependent RNA helicase SUPV3L1 |
| TRIAE_CS42_2DS_TGACv1_178212_AA0592790 | SAC3/GANP/Nin1/mts3/eIF-3 p25 |
| TRIAE_CS42_2DL_TGACv1_160098_AA0546700 | Myb/SANT-like DNA-binding protein |
| TRIAE_CS42_2DS_TGACv1_177707_AA0582780 | pfkB-like carbohydrate kinase family protein |
| TRIAE_CS42_2DS_TGACv1_178855_AA0601830 | 4-coumarate:CoA ligase 2 |
| TRIAE_CS42_2DL_TGACv1_159638_AA0540810 | Unknown |
| TRIAE_CS42_2DL_TGACv1_158120_AA0509790 | Peroxidase |
| TRIAE_CS42_2DL_TGACv1_158042_AA0507110 | Cysteine/Histidine-rich C1 domain family protei |
| TRIAE_CS42_2DS_TGACv1_177392_AA0575450 | Receptor protein kinase, putative |
| TRIAE_CS42_2DS_TGACv1_177277_AA0571960 | 3'-N-debenzoyl-2'-deoxytaxol N-benzoyltransfe |
| TRIAE_CS42_2DL_TGACv1_158042_AA0507140 | Leucine-rich repeat receptor-like protein kinase |
| TRIAE_CS42_2DS_TGACv1_177747_AA0583750 | Protein kinase |
| TRIAE_CS42_2DL_TGACv1_158085_AA0508400 | Protein of unknown function (DUF581) |
| TRIAE_CS42_2DL_TGACv1_159259_AA0535530 | Wound-induced protein |
| TRIAE_CS42_2DS_TGACv1_177577_AA0580340 | P-loop containing nucleoside triphosphate hydr |
| TRIAE_CS42_2DS_TGACv1_178278_AA0593740 | Cytochrome P450 |
| n/a | - |
| TRIAE_CS42_2DS_TGACv1_178414_AA0595260 | WUSCHEL-related homeobox 11 |
| TRIAE_CS42_2DS_TGACv1_178429_AA0595420 | Trehalose 6-phosphate phosphatase |
| TRIAE_CS42_2DL_TGACv1_158388_AA0517230 | Germin-like protein 4-1 |
| TRIAE_CS42_2DL_TGACv1_158637_AA0523690 | AAA domain UvrD/REP helicase N-terminal dom |
| Kelch Motif | - |
| TRIAE_CS42_2DS_TGACv1_177249_AA0570400 | Putative glutathione S-transferase |
| TRIAE_CS42_2DS_TGACv1_178283_AA0593780 | Unknown |
| TRIAE_CS42_2DL_TGACv1_158175_AA0511290 | Germin-like protein 4-1 |
| n/a | - |
| TRIAE_CS42_2DL_TGACv1_159768_AA0542480 | Unknown |
| n/a | - |
| TRIAE_CS42_2DS_TGACv1_177302_AA0572660 | 2-oxoglutarate (2OG) and Fe(II)-dependent oxy |

| | |
|---|---|
| TRIAE_CS42_2DL_TGACv1_158354_AA0516480 | Peroxidase |
| TRIAE_CS42_2DL_TGACv1_161020_AA0556020 | Transcription cofactor |
| TRIAE_CS42_2DL_TGACv1_160112_AA0546950 | Wound-responsive family protein |
| TRIAE_CS42_2DL_TGACv1_159072_AA0531800 | Putative receptor like protein kinase |
| TRIAE_CS42_2DS_TGACv1_177373_AA0574940 | Zinc-finger protein |
| TRIAE_CS42_2DS_TGACv1_177302_AA0572740 | Disease resistance protein RPM1 |
| TRIAE_CS42_2DS_TGACv1_177503_AA0578970 | Unknown |
| TRIAE_CS42_2DS_TGACv1_177679_AA0582290 | Lectin-domain containing receptor kinase |
| TRIAE_CS42_2DL_TGACv1_159581_AA0540110 | Heavy metal transport/detoxification superfam |
| TRIAE_CS42_2DS_TGACv1_178829_AA0601440 | Glucose-6-phosphate/phosphate translocator 2 |
| TRIAE_CS42_2DL_TGACv1_161137_AA0557010 | RING-H2 finger protein |
| TRIAE_CS42_2DL_TGACv1_158540_AA0521470 | Peroxidase |
| TRIAE_CS42_2DL_TGACv1_158303_AA0515200 | Brassinosteroid signalling positive regulator (BZ |
| TRIAE_CS42_2DS_TGACv1_178325_AA0594150 | FAD-binding Berberine |
| TRIAE_CS42_2DL_TGACv1_157936_AA0502560 | 2-oxoglutarate (2OG) and Fe(II)-dependent oxy |
| TRIAE_CS42_2DL_TGACv1_157936_AA0502570 | 2-oxoglutarate (2OG) and Fe(II)-dependent oxy |
| TRIAE_CS42_2DL_TGACv1_159181_AA0533940 | Peroxidase |
| TRIAE_CS42_2DL_TGACv1_158070_AA0507860 | Cytochrome P450 |
| n/a | - |
| TRIAE_CS42_2DL_TGACv1_162095_AA0561750 | Quinone oxidoreductase |
| n/a | - |
| n/a | - |
| TRIAE_CS42_2DS_TGACv1_178484_AA0596180 | WRKY transcription factor 46 |
| TRIAE_CS42_2DL_TGACv1_157927_AA0502330 | Peroxidase |
| TRIAE_CS42_2DL_TGACv1_162184_AA0562070 | Desiccation-related protein PCC13-62 |
| TRIAE_CS42_2DS_TGACv1_178030_AA0589590 | Quinone oxidoreductase |
| n/a | - |
| TRIAE_CS42_2DS_TGACv1_177265_AA0571140 | Potassium transporter |
| TRIAE_CS42_2DS_TGACv1_177631_AA0581600 | Peroxidase |
| n/a | - |
| TRIAE_CS42_2DL_TGACv1_158553_AA0521690 | Pectinesterase inhibitor domain containing pro |
| TRIAE_CS42_2DS_TGACv1_177894_AA0586810 | Jasmonate-zim-domain protein 1 |
| n/a | - |
| TRIAE_CS42_2DL_TGACv1_160467_AA0550820 | Thaumatin-like protein |
| TRIAE_CS42_2DS_TGACv1_178097_AA0590730 | Peroxidase |
| TRIAE_CS42_2DS_TGACv1_177232_AA0569640 | Dirigent-like protein |
| TRIAE_CS42_2DL_TGACv1_160265_AA0548860 | Unknown |
| TRIAE_CS42_2DS_TGACv1_177444_AA0577250 | Unknown |
| n/a | Nuclear pore complex scaffold, nucleoporin |
| TRIAE_CS42_2DL_TGACv1_157922_AA0501820 | Protease inhibitor/seed storage/lipid transfer f |
| TRIAE_CS42_2DL_TGACv1_157948_AA0503070 | Glutaredoxin |
| TRIAE_CS42_2DS_TGACv1_178216_AA0592850 | Retrotransposon protein, putative, Ty3-gypsy s |
| n/a | - |
| TRIAE_CS42_2DL_TGACv1_158819_AA0526620 | WRKY transcription factor 32 |
| TRIAE_CS42_2DL_TGACv1_158826_AA0526780 | Basic helix-loop-helix (BHLH) Transcription Fact |
| n/a | - |

| | |
|---|---|
| TRIAE_CS42_2DL_TGACv1_160663_AA0552860 | BAG family molecular chaperone regulator 3 |
| n/a | - |
| TRIAE_CS42_2DL_TGACv1_159135_AA0533070 | Unknown |
| TRIAE_CS42_2DL_TGACv1_158211_AA0512570 | Peroxidase |
| TRIAE_CS42_2DL_TGACv1_159535_AA0539490 | Protease inhibitor/seed storage/lipid transfer p |
| TRIAE_CS42_2DS_TGACv1_179053_AA0603910 | Chemocyanin |
| TRIAE_CS42_2DS_TGACv1_179236_AA0605510 | WRKY transcription factor 4 |
| n/a | Unknown |
| TRIAE_CS42_2DL_TGACv1_160694_AA0553200 | Nitrate transporter 1.2 (NPF4) |
| TRIAE_CS42_2DL_TGACv1_160902_AA0555040 | Heavy metal transport/detoxification superfam |
| TRIAE_CS42_2DS_TGACv1_179382_AA0606710 | Dihydroflavonol-4-reductase |
| TRIAE_CS42_2DL_TGACv1_157948_AA0503060 | NAC transcription factor |
| TRIAE_CS42_2DS_TGACv1_177143_AA0566600 | NBS-LRR disease resistance protein family-3 |
| n/a | - |
| TRIAE_CS42_2DL_TGACv1_160875_AA0554830 | Unknown |
| TRIAE_CS42_2DL_TGACv1_160902_AA0555030 | Heavy metal transport/detoxification superfam |
| TRIAE_CS42_2DL_TGACv1_162805_AA0563610 | Anthocyanidin reductase |
| n/a | - |
| n/a | - |
| n/a | - |
| n/a | - |
| n/a | - |

| TGAC contig start position in I' | IWGSC RefSeqv1 Gene Name |
|---|---|
| 2D: 1587841 | TraesCS2D02G001700 |
| 2D: 498767044 | TraesCS2D02G391500 |
| 2D: 403496642 | TraesCS2D02G408400LC |
| 2D: 481988679 | TraesCS2D02G377600 |
| 2D: 402938727 | TraesCS2D02G313300 |
| 2D: 162226243 | - |
| 2D: 288139267 | - |
| 2D: 127085359 | TraesCS2D02G182800 |
| 2D: 448689745 | TraesCS2D02G350700 |
| 2D: 14088978 | - |
| 2D: 302250516 | TraesCS2D02G252100 |
| 2D: 508405693 | - |
| 2D: 559295489 | TraesCS2D02G558400LC |
| 2D: 141865595 | - |
| 2D: 141868923 | - |
| 2D: 2466831 | TraesCS2D02G003600LC |
| 2D:106351711 | - |
| 2D: 104909076 | - |
| 2D: 55270733 | - |
| 2D: 477269807 | TraesCS2D02G373900 |
| 2D: 141863644 | - |
| 2D: 169624953 | - |
| 2D: 131279750 | TraesCS2D02G187400 |
| 2D: 87258980 | TraesCS2D02G145300 |
| 2D: 2088871 | - |
| 2D: 425591092 | - |
| 2D: 260016412 | - |
| 2D: 12893803 | - |
| 2D: 614726221 | - |
| 2D: 304088006 | - |
| 2D: 420194171 | - |
| 2D: 11558528 | TraesCS2D02G028200 |
| 2D: 11574620 | TraesCS2D02G028300 |
| 2D: 12877640 | - |
| 2D: 12873839 | - |
| 2D: 529008008 | TraesCS2D02G415100 |
| 2D: 528994683 | TraesCS2D02G414900 |
| 2D: 12660346 | TraesCS2D02G031700 |
| 2D: 439813636 | - |
| 2D: 112923076 | - |
| 2D: 595165983 | TraesCS2D02G500600 |
| 2D: 185187419 | - |
| 2D: 93263223 | - |
| 2D: 310225990 | - |

| | |
|---|---|
| 2D: 86526289 | TraesCS2D02G145100 |
| 2D: 598305972 | TraesCS2D02G618300LC |
| 2D: 472408832 | - |
| 2D: 12550693 | TraesCS2D02G030800 |
| 2D: 43227664 | TraesCS2D02G096400LC |
| 2D: 439803307 | - |
| 2D: 86717283 | TraesCS2D02G145200 |
| 2D: 477259990 | TraesCS2D02G373800 |
| 2D: 162229763 | - |
| 2D: 446256266 | TraesCS2D02G348200 |
| 2D: 126114348 | - |
| 2D: 327843578 | - |
| 2D: 184849390 | - |
| 2D: 178352970 | - |
| 2D: 601215188 | TraesCS2D02G507800 |
| 2D: 491402626 | - |
| 2D: 573845078 | TraesCS2D02G579200LC |
| 2D: 416757383 | TraesCS2D02G323600 |
| 2D: 86076159 | TraesCS2D02G144800 |
| 2D: 621363664 | - |
| 2D: 29675518 | - |
| 2D: 92921251-92919093 | TraesCS2D02G150400 |
| 3B: 525631539-525630689 | - |
| 2D: 601984319-601982942 | TraesCS2D02G509700 |
| 2D: 440591045-440594173 | TraesCS2D02G344400 |
| 2D: 5388743-5392838 | TraesCS2D02G010900 |
| 2D: 2614280-2615365 | - |
| 2D: 440633771-440634803 | TraesCS2D02G344600 |
| 2D: 29880394-29875373 | TraesCS2D02G070800 |
| 2D: 562340880-562341688 | TraesCS2D02G453400 |
| 2D: 627363415-627360988 | TraesCS2D02G552100 |
| 2D: 195421681-195423551 | TraesCS2D02G227000 |
| 2D: 11964758-11962394 | TraesCS2D02G029300 |
| 2D: 179350119-179349741 | - |
| 2D: 52229894-52227196 | TraesCS2D02G100200 |
| 2D: 112102472-112099168 | TraesCS2D02G168200 |
| 2D: 566835044-566837828 | TraesCS2D02G460100 |
| 2D: 551861960-551876222 | TraesCS2D02G441300 |
| 2D: 434532553-434532173 | Homoeolog of TraesCS2A02G111200 |
| 2D: 192186524-192185419 | TraesCS2D02G224700 |
| 2D: 233436061-233430328 | Homoeolog of TraesCS2B02G126600 |
| 2D: 566979655-566981551 | TraesCS2D02G460200 |
| 2D: 413275323-413273542 | - |
| 2D: 448502203-448500805 | - |
| 2D: 133481813-133479344 | - |
| 2D: 11459670-11462723 | TraesCS2D02G027600 |

| | |
|---|---|
| 2D: 643641622-643640294 | TraesCS2D02G583200 |
| 2D: 648170503-648165702 | TraesCS2D02G594100 |
| 2D: 587106893-587107591 | TraesCS2D02G487000 |
| 2D: 472466848-472469314 | TraesCS2D02G367900 |
| 2D: 38183834-38184683 | TraesCS2D02G088100 |
| 2D: 11549999-11555148 | TraesCS2D02G028100 |
| 2D: 125508603-125509399 | TraesCS2D02G197200LC |
| 2D: 75001397-74998859 | TraesCS2D02G129100 |
| 2D: 423212613-423213346 | TraesCS2D02G330200 |
| 2D: 194189950-194187604 | TraesCS2D02G226000 |
| 2D: 563959601-563958427 | TraesCS2D02G455800 |
| 2D: 635760657-635758961 | TraesCS2D02G566800 |
| 2D: 286986962-286984334 | - |
| 2D: 73452287-73454525 | TraesCS2D02G126400 |
| 2D: 500793696-500791572 | TraesCS2D02G392900 |
| 2D: 500694513-500688336 | TraesCS2D02G392800 |
| 2D: 644016313-644017780 | TraesCS2D02G719900LC |
| 2D: 589629928-589627845 | TraesCS2D02G491000 |
| 2D: 593229385-593226612 | - |
| 2D: 323516017-323517158 | TraesCS2D02G264400 |
| 2D: 465954181-465954613 | - |
| 2D: 155475292-155477882 | - |
| 2D: 145888093-145893104 | TraesCS2D02G198100 |
| 2D: 643901039-643902572 | TraesCS2D02G584700 |
| 2D: 370175621-370176979 | TraesCS2D02G288700 |
| 2D: 73820169-73819158 | TraesCS2D02G127000 |
| 2D: 422881740-422883049 | - |
| 2D: 58781534-58776915 | TraesCS2D02G106600 |
| 2D: 60137542-60139234 | TraesCS2D02G108500 |
| 2D: 35003316-34999870 | - |
| 2D: 618700523-618701353 | TraesCS2D02G536100 |
| 2D: 120256719-120254261 | TraesCS2D02G176800 |
| 2D: 420521791-420525990 | - |
| Un: 162316211-162315699 | - |
| 2D: 59728494-59727019 | TraesCS2D02G107600 |
| 2D: 18187313-18188975 | TraesCS2D02G042600LC |
| 2D: 405409868-405412352 | TraesCS2D02G316100 |
| 2D: 186959906-186957508 | TraesCS2D02G245700LC |
| 2D: 261784439-261784054 | Similar to TraesCS6A02G175000 |
| 2D: 377151112-377150306 | TraesCS2D02G295000 |
| 2D: 466683167-466684403 | TraesCS2D02G361400 |
| 2D: 109901632-109909785 | TraesCS2D02G165800 |
| 2D: 650859941-650858111 | - |
| 2D: 588677697-588676060 | TraesCS2D02G489700 |
| 2D: 333498415-333503448 | TraesCS2D02G270000 |
| 2D: 29743490-29746782 | - |

| | |
|---|---|
| 2D: 435198153-435196765 | TraesCS2D02G340900 |
| 2D: 125343657-125344934 | - |
| 2D: 632454827-632457525 | - |
| 2D: 602711264-602712722 | TraesCS2D02G511200 |
| 2D: 547775108-547774264 | TraesCS2D02G438200 |
| 2D: 95417890-95417172 | TraesCS2D02G152100 |
| 2D: 56256125-56253778 | TraesCS2D02G104500 |
| 2D: 533173295-533169963 | Similar to TraesCS4B02G057100 |
| 2D: 446306205-446312471 | TraesCS2D02G348400 |
| 2D: 620159918-620160800 | - |
| 2D: 125733989-125729785 | TraesCS2D02G181300 |
| 2D: 466682639-466685866 | TraesCS2D02G361400 |
| 2D: 14909653-14910981 | - |
| 2D: 620223828-620225080 | - |
| 2D: 614826542-614825803 | - |
| 2D: 620153338-620154529 | TraesCS2D02G540000 |
| 2D: 584613732-584606715 | TraesCS2D02G481900 |
| 2D: 11,291,329-11,291,519 | - |
| 2D: 641,484,219-641,484,862 | - |
| 2D: 11,549,905-11,550,092 | - |
| 2D: 16,346,791-16,346,260 | - |
| 2D: 12,555,127-12,555,463 | - |
