## Supplementary material for "Identification of QTL and Underlying Genes for Root System Architecture associated with Nitrate Nutrition in Hexaploid Wheat": Table S3

**Table S3.** Alignment statistics after using HISAT2 to align the RNA-seq samples to the Chinese Spring reference genomes.

| Group | Sample | Number of Sequencing reads | Number of sequencing reads aligned (1^st^ pass) | Number of sequencing reads aligned uniquely | Number of sequencing reads aligned uniquely after remove duplicates | Bp mapped | Sequencing reads aligned uniquely after remove duplicates (to unmapped read assembly) | Bp mapped (of unmapped read assembly) |
| --- | --- | --- | --- | --- | --- | --- | --- | --- |
| A | 2D1 | 39,810,948 | 35,095,839 | 30,462,674 | 23,988,769 | 258,375,578 | 229,029 | 6,127,012 |
| A | 2D2 | 33,218,456 | 26,721,116 | 23,375,577 | 18,719,507 | 251,797,502 | 250,793 | 6,399,364 |
| A | 2D3R | 45,922,198 | 39,890,448 | 34,655,561 | 26,812,582 | 261,430,002 | 236,496 | 5,700,772 |
| A | 2D7R | 52,047,398 | 46,578,833 | 40,526,437 | 32,058,907 | 277,941,174 | 309,032 | 6,947,487 |
| B | 2D4R | 40,822,742 | 36,421,999 | 31,311,152 | 25,346,164 | 267,226,179 | 193,180 | 4,562,473 |
| B | 2D5 | 41,214,782 | 36,547,602 | 31,648,643 | 24,190,602 | 262,069,625 | 207,809 | 5,541,350 |
| B | 2D6 | 39,469,064 | 34,326,561 | 29,438,683 | 22,274,621 | 257,019,220 | 161,092 | 4,582,661 |
| B | 2D8 | 47,363,162 | 42,365,355 | 36,662,284 | 29,147,092 | 280,891,736 | 228,139 | 5,413,053 |
