## Supplementary material for "Identification of QTL and Underlying Genes for Root System Architecture associated with Nitrate Nutrition in Hexaploid Wheat": Table S5

**Table S5.** Composition of ¼ Hoagland’s nutrient solution used in wheat seedling study.

| For low N treatment, Ca(NO_3_)_2_·4H_2_O and KNO_3_ were removed and replaced with 0.7 mM CaSo_4_·½H_2_O and 1.5 mM KCl. | | | |
| --- | --- | --- | --- |
| Macronutrients | mM | Micronutrients | µM |
| (NH_4_)_3_PO_4_ | 0.25 | CuSO_4_·5H_2_O | 3.0 |
| Ca(NO_3_)_2_·4H_2_O | 0.7 | MnCl_2_·4H_2_O | 63.05 |
| MgSO_4_·7H_2_O | 1.02 | MoO_3_ | 1.39 |
| KNO_3_ | 1.5 | ZnSO_4_·7H_2_O | 8.0 |
| H_3_BO_3_ | 0.46 | FeHEDTA | 77.0 |
