## Supplementary material for "Identification of QTL and Underlying Genes for Root System Architecture associated with Nitrate Nutrition in Hexaploid Wheat": Figure S2

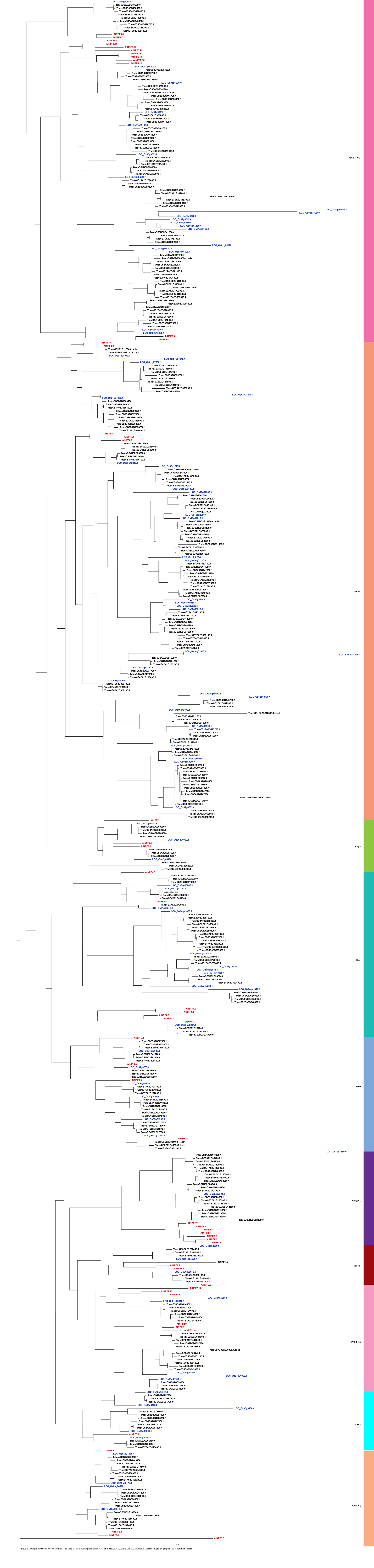


Fig. S2. Phylogenetic tree of protein families comparing the NPF family protein sequences of *A. thaliana*, *O. sativa* L and *T. aestivum* L. Branch lengths are proportional to substitution rate.
