## Supplementary material for "Identification of QTL and Underlying Genes for Root System Architecture associated with Nitrate Nutrition in Hexaploid Wheat": Data S1 and S2: Data S1.ijm

```
// Color Thresholder 2.0.0-rc-43/1.50e
// Autogenerated macro, single images only!
dir = getDirectory("image"); 
min=newArray(3);
max=newArray(3);
filter=newArray(3);
a=getTitle();
//makeRectangle(892, 784, 1836, 1520);
//run("Crop");
run("HSB Stack");
run("Convert Stack to Images");
selectWindow("Hue");
rename("0");
selectWindow("Saturation");
rename("1");
selectWindow("Brightness");
rename("2");
min[0]=0;
max[0]=120;
filter[0]="pass";
min[1]=0;
max[1]=255;
filter[1]="pass";
min[2]=14;
max[2]=255;
filter[2]="pass";
for (i=0;i<3;i++){
  selectWindow(""+i);
  setThreshold(min[i], max[i]);
  run("Convert to Mask");
  if (filter[i]=="stop")  run("Invert");
}
imageCalculator("AND create", "0","1");
imageCalculator("AND create", "Result of 0","2");
for (i=0;i<3;i++){
  selectWindow(""+i);
  close();
}
selectWindow("Result of 0");
close();
selectWindow("Result of Result of 0");
rename(a);
run("Analyze Particles...", "size=1000-Infinity circularity=0.00-0.6 show=[Overlay] display in_situ");
index = lastIndexOf(a, "."); 
if (index!=-1) a = substring(a, 0, index);
b = a;
a = a + ".csv"; 
saveAs("Measurements", dir+a);
//selectWindow(b + ".jpg");
//saveAs("PNG",dir+b);
//close();  
```
