## Supplementary material for "Identification of QTL and Underlying Genes for Root System Architecture associated with Nitrate Nutrition in Hexaploid Wheat": Data S1 and S2: Data S2.R

```
### Examples will use 'phenotype' in place of trait name for each line
### Make sure genotype column is called ‘Line’ before importing
### Make sure experiment number column is named ‘Experiment.Number’
### data file renamed to root_data using:

root_data <- file name

install.packages('lme4')
library('lme4')

#########################
### To calculate BLUEs###
#########################
model1 <- lm(phenotype ~ Line + Experiment.Number, data=root_data)
sum1 <- summary(model1)

estimates <- sum1$coefficients[,'Estimate']
BLUEs <- estimates
BLUEs[2:length(BLUEs)] <- BLUEs[2:length(BLUEs)]+BLUEs[1]
write.csv(BLUEs, file = "BLUEs.csv")

### To remove all & restart
rm(BLUEs, estimates, model1, sum1)

#########################
### To calculate BLUPs###
#########################
model3<- lmer(phenotype ~ Line + (1|Experiment.Number), data=root_data)
BLUPs <-fixef(model3)
write.csv(BLUPs, file = "BLUPs.csv")
```
